# ABI3 regulates ABI1 function to control cell length in primary root elongation zone

**DOI:** 10.1101/2023.10.16.562476

**Authors:** Saptarshi Datta, Drishti Mandal, Sicon Mitra, Ronita Nag Chaudhuri

## Abstract

Post-embryonic primary root growth is effectively an interplay of several hormone signalling pathways. Here, we show that the ABA-responsive transcription factor ABI3, controls primary root growth through regulation of JA signalling molecule *JAZ1* along with ABA responsive factor ABI1. In absence of ABI3, primary root elongation zone is shortened with significantly reduced cell length. Expression analyses and ChIP based assays indicate that ABI3 negatively regulates *JAZ1* expression by occupying its upstream regulatory sequence and enriching repressive histone modification mark H3K27 trimethylation, thereby occluding RNAPII occupancy. Previous studies have shown that JAZ1 interacts with ABI1, the protein phosphatase 2C, that works during ABA signalling. Our results indicate that in absence of ABI3, when *JAZ1* expression levels are high, ABI1 protein shows increased stability, compared to when JAZ1 is absent, or ABI3 is overexpressed. Consequently, in *abi3* mutant, due to higher stability of ABI1, reduced phosphorylation of plasma membrane H^+^ATPase (AHA2) occurs. HPTS staining further indicated that, *abi3* root cell apoplasts show reduced protonation, compared to wild type and ABI3 overexpressing seedlings. Such impeded proton extrusion, negatively affects cell length in primary root elongation zone. ABI3 therefore controls cell elongation in primary root by affecting ABI1-dependent protonation of root cell apoplasts. In summary, ABI3 controls expression of JAZ1 and in turn modulates function of ABI1 to regulate cell length in the elongation zone during primary root growth.

## Introduction

Root system is an underground plant organ that primarily functions towards uptake and translocation of water and essential nutrients, along with providing the plant its necessary anchorage. Primary root and lateral root are the two main root types that make up the root system architecture. Multilayered hormonal crosstalk dictates effective root development based on prevailing environmental cues. Primary root growth is indeterminate and is critically regulated by two developmental mechanisms, cell division in the root apical meristem (RAM) and cell elongation in the elongation zone (EZ). Several phytohormones including auxin, cytokinin, ethylene, abscisic acid (ABA), gibberellic acid (GA) and jasmonic acid (JA) have been shown to play major roles in regulating the epidermal cell length in the elongation zone (Barbez et al., 2017, Růzicka et al., 2007, Chen et al., 2011, Shani et al., 2013, Street et al., 2016, Miao et al., 2021). Auxin serves as the pivotal player around which other plant hormones converge to coordinate regulation of root development.

ABA is a vital stress hormone that is also known to regulate several developmental processes such as seed germination, controlling stomatal aperture and seedling growth (Finkelstein et al., 2008, Fujita et al., 2011, Hauser et al., 2011, Nakashima and Yamaguchi-Shinozaki, 2013). It is also involved in modulating root system architecture by inhibiting lateral root development, regulating primary root growth in a dose-dependent manner and maintaining root hydrotropism (De Smet et al., 2003, Dietrich et al., 2017, Luo et al., 2023, Sun et al., 2018). Over the past few decades, significant advancements have been achieved in comprehending ABA signalling (Ali et al., 2020, Hubbard et al., 2010, Raghavendra et al., 2010). ABA is perceived by the pyrabactin resistance 1 (PYR1)/ PYR1-like (PYL)/ regulatory components of the ABA receptor (RCAR) protein family, which leads to recruitment of the clade A protein phosphatases type 2C (PP2Cs), such as ABA-insensitive 1 (ABI1), ABI2, hypersensitive to ABA1 (HAB1) and PP2CA to form a PYR/RCAR-PP2C complex and inhibit phosphatase activity (Antoni et al., 2012, Ma et al., 2009, Nishimura et al., 2009, Nishimura et al., 2010). Thereafter, Sucrose non– fermenting1–related protein kinase 2 (SnRK2s) are relieved from PP2C-SnRK2 complexes that phosphorylate downstream transcription factors such as ABI3, ABI4 and ABI5 to activate ABA responsive element (ABRE) driven downstream gene expression (Fujii et al., 2007, Fujii and Zhu, 2009, Nakashima et al., 2009). In absence of ABA, the interaction between PP2Cs and PYR/PYL/RCARs is blocked, resulting in PP2C activation. Consequently, PP2Cs interact and impede the kinase activity of SnRK2s by dephosphorylating the kinase activation loop, thereby suppressing ABA signalling (Ma et al., 2009, Melcher et al., 2009, Miyazono et al., 2009, Nishimura et al., 2009, Park et al., 2009). The co-receptor ABI1 plays a dual role; it functions as a component in a negative feedback mechanism within the ABA signalling pathway, while also serving as a central point for crosstalk with other hormones and undergoes regulation at multiple levels (Julian et al., 2019, Kong et al., 2015, Tajdel et al., 2016). In regulating primary root growth, ABA signalling works in biphasic manner. At low concentrations such as 0.1 µM, ABA stimulates primary root growth; however, at higher concentrations such as 10 µM, it impedes primary root growth (Miao et al., 2021, Sun et al., 2018). Additionally, ABI1 has been shown to interact with the C-terminus of the plasma membrane H^+^-ATPase (AHA2) to form a complex and dephosphorylates it at Thr^947^, thereby suppressing H^+^ efflux. Under low ABA concentration cytosolic ABA receptors bind to PP2Cs, rendering AHA2 active, causing apoplastic H^+^ extrusion that effectively results in cell elongation at the EZ of primary root (Miao et al., 2021).

JA is a well-known lipid-derived phytohormone that regulates plant growth and development, as well as response to stress. JA governs multiple aspects of root growth, including inhibition of primary root growth (Chen et al., 2011), regulation of lateral root formation and root regeneration (Cai et al., 2014, Ye et al., 2020, Zhang et al., 2019). Jasmonate limits primary root elongation by diminishing both cell proliferation in the meristematic zone and cell size in the elongation zone, indicating that the regulation is complex, encompassing numerous factors (Chen et al., 2011, Shukla et al., 2020, Staswick et al., 1992). JA is perceived by F box protein CORONATINE INSENSITIVE1 (SCF^COI1^ complex) and targets JASMONATE ZIM DOMAIN (JAZ) proteins for degradation which are negative regulators of the JA signalling (Chini et al., 2007, Thines et al., 2007, Xu et al., 2002, Yan et al., 2009). Degradation of JAZ repressors mitigate the suppression of downstream transcription factors such as MYC2,3,4 and activates JA signalling (Chini et al., 2007, Pauwels and Goossens, 2011, Thines et al., 2007). Previous studies have demonstrated JAZ proteins interact with multiple transcription factors, thereby affecting intricate interplay between JA and other plant hormones. JAZ1, JAZ3 and JAZ9 physically interact with EIN3/EIL1 to mediate the crosstalk between JAZs and ethylene (Zhu et al., 2011). Additionally, JAZs also interact with DELLA, a negative regulator of GA signalling (Hou et al., 2013). Recently, physical interactions have been reported for JAZ1, JAZ5 and JAZ8 with ABI3, suggesting the existence of a cross-talk between JAZs and ABA signaling (Pan et al., 2020). Interestingly, it has been shown that auxin can stimulate *JAZ1* expression in the primary root (Grunewald et al., 2009). It is therefore relevant to assert that multifaceted crosstalk exists between JAZs and various other phytohormones.

Multiple studies have presented evidence supporting the existence of interplay between ABA and JA signalling. ABA is known to promote both ABA and JA biosynthesis in Arabidopsis, while this phenomenon is reduced in *coi1* mutant and quadruple JAZ mutant *jazQ* (Campos et al., 2016) (Ju et al., 2019, Pan et al., 2020). Moreover, *pyl4* and *pyl5* knockout mutants show hypersensitivity to JA indicating that PYR/PYL/RCAR receptors function in the crosstalk between JA and ABA signalling (Lackman et al., 2011). On the other hand, MYC2 overexpressing plants have been shown to be hypersensitive to ABA (Abe et al., 2003). Further studies have demonstrated a direct interaction between PYL6 and MYC2. Interestingly, the *pyl6* mutant displayed heightened sensitivity to the combined presence of JA and ABA compared to ABA alone (Aleman et al., 2016). Moreover, it is known that the JA and ABA regulate the process of stomatal closure in Arabidopsis by modulating the protein kinase known as OPEN STOMATA1 (Yin et al., 2016). These studies taken together implicate a link between ABA and JA signalling.

While the interplay between the ABA and JA signalling has been previously documented, there remains limited understanding regarding the specific impact of their crosstalk during primary root development. In this study, we have investigated the molecular basis of coordination between ABA signalling and *JAZ1* during primary root growth in Arabidopsis. We show that ABI3 regulates primary root elongation by negatively regulating *JAZ1.* Absence of ABI3 and presence of JAZ1 promotes ABI1 protein stability that leads to reduced phosphorylation of AHA2 (PM H^+^-ATPase). This effectively decreased protonation of root cell apoplast. Such alkalinization of the apoplast led to shortened cell length and thus reduced primary root elongation. Our findings uncover a novel paradigm underlying the interaction between JA and ABA signalling during regulation of primary root growth in Arabidopsis.

## Materials and Methods

For a more detailed description of the Materials and Methods see Method S1 section.

### Plant materials and growth condition

All genotypes utilized in this study are in the *Arabidopsis thaliana* ecotype Columbia (*Col-0*) background. The following Arabidopsis seed stocks were used: *abi3* (was kindly provided by Prof. Eiji Nambara, University of Toronto, Canada) (Bedi et al., 2016, Nambara et al., 1994), *abi1* (NASC stock ID N538866), *jaz1* (NASC stock ID N2107784) (Shukla et al., 2020). For stable line generation, 35S::ABI3 (*ABI3OX-1, ABI3OX-2*), proJAZ1::GFP and proABI3::GFP constructs were introduced into *Col-0* plants by the Agrobacterium-mediated floral dip method The proJAZ1::GFP construct was also introduced into *abi3* mutant plants. The mutant lines were confirmed by PCR, using primers as listed in Table S1. The seeds were surface sterilized (Bedi and Nag Chaudhuri, 2018, Bedi et al., 2016, Mandal et al., 2023, Sengupta and Nag Chaudhuri, 2020) and vernalized at 4°C for 2 days in darkness, and then plated on Murashige and Skoog (MS) medium. For morphological analyses, seedlings were grown vertically (22°C, 16 h light/8 h dark) in MS medium for the indicated number of days. *Nicotiana benthamiana* seeds were directly sown on potting soil (Soilrite Mix, Keltech Energies, India) and grown at 25°C in the plant growth room.

### Cloning and protein purification

For full length coding sequences (CDS) of *ABI3* (2870 bp) and for promoter::GUS/GFP reporter fusion construct (proJAZ1::GUS/GFP – 667 bp) sequences of the genes were amplified and was first cloned into pENTR/d-TOPO (Invitrogen, Life Technologies). For over-expression under *CaMV 35S* promoter (35S::ABI3) and for proJAZ1::GUS/GFP reporter fusion, entry clones were recombined with pK7WG2D and pKGWFS7 respectively using LR Clonase II (Thermo Fisher Scientific). proABI3::GUS/GFP (1.6 kb) construct was previously generated (Bedi and Nag Chaudhuri, 2018). All the recombinant plasmids were subsequently transformed into *Agrobacterium tumefaciens* strain LBA4404.

Full length (CDS) clone of ABI1 (1302 bp) and ARR1 (2070 bp) was generated using cDNA of *Col-0* as template and ligated into T-vector pMD20 (Takara Bio, Japan). All the primers used for cloning were listed in Table S1. Positive clones were subcloned into pET32A and transformed into *E.coli* BL21 (DE3) plysS. The cells were induced with 0.5 mM IPTG at 28°C and 22°C overnight for ABI1_myc and ARR1_His proteins respectively. The cells were then harvested in lysis buffer (50 mM NaH_2_PO_4_, 300 mM NaCl, 10 mM imidazole) and subjected to sonication. The lysate was centrifuged and the soluble cell fraction was incubated with Ni^2+^ –NTA (Qiagen) beads for 3 hr. After washing the beads with wash buffers (50 mM NaH_2_PO_4_, 300 mM NaCl) containing 10, 30 and 50 mM imidazole ABI1_myc and ARR1_His proteins were finally eluted with elution buffer (50 mM NaH_2_PO_4_, 300 mM NaCl, 250 mM Imidazole).

### Measurement of root length and EZ cell length

For morphological analysis, root lengths were measured from root–hypocotyl junction to root tip using ImageJ software (Mandal et al., 2023). Elongation zone (EZ) length and epidermal cell length of EZ were measured as described (Růzicka et al., 2007, Strader et al., 2010, Verbelen et al., 2006) using ImageJ software. At least 20 seedlings of each genotype were measured and three independent biological replicates were done.

### Staining and microscopic analyses

To visualize EZ, roots were dipped in propidium iodide solution (10 µg/ml) for 30 sec to stain the cell wall and were analysed under Leica SP8 confocal microscope (excitation 535 nm and emission 610–640 nm) (Mandal et al., 2023). For *ABI3* and *JAZ1* localization, stable transgenic lines were visualized under Leica fluorescence microscope DM4B equipped with Leica DFC3000 camera with GFP filter settings.

For 8-hydroxypyrene-1,3,6-trisulfonic acid trisodium salt (HPTS) staining, 5-day old seedlings were incubated in ½ MS media supplemented with 1 mM HPTS (Barbez et al., 2017). The roots were then mounted in the same growth media on a microscopic slide. Imaging was performed using Leica SP8 confocal microscope equipped with hybrid laser detectors. Fluorescent signals for the protonated HPTS form (excitation 405 nm, emission peak 514 nm), as well as the deprotonated HPTS form (excitation 448 nm, emission peak 514 nm) were detected. Image analysis and fluorescent quantification was done using ImageJ software.

### RNA isolation for gene expression analyses

Roots of WT, *abi3*, *ABI3OX-1* and *ABI3OX-2* were harvested at the indicated number of days. Detailed methods are provided in Method S1 section.

### Chromatin immunoprecipitation assay

Nuclei isolation and subsequent ChIP assay with anti-ABI3 (BioBharati Life Sciences, Kolkata West Bengal, India) (Bedi et al., 2016, Sengupta et al., 2020), anti-RNAPII (Abcam ab817), anti-H3K4me3 (Abcam ab8580), anti-H3K27me3 (Abcam ab195477) and anti-H3 (BioBharati Life Sciences, India) antibody were performed as described previously (Bedi et al., 2016, Mandal et al., 2023, Sengupta et al., 2020). Detailed methods are provided in Method S1 section.

### Real-time PCR and data analyses

Real-time PCR and subsequent data analyses was done as described previously (Mandal et al., 2023). Detailed methods of real-time PCR and data analyses are provided in Method S1 section.

### Fluorometric *GUS* assay

Fluorometric GUS assay was performed as described previously (Bedi and Nag Chaudhuri, 2018, Mandal et al., 2023, Sengupta et al., 2020). Comprehensive methodologies are given in the Method S1 section.

### Histochemical GUS staining

GUS staining of the agrobacterium infiltrated Nicotiana leaves were done according to (Fang et al., 2015). Detailed description is given in the Method S1 section.

### Total protein isolation

Detailed description of total protein isolation from WT and *abi3* seedlings are given in the Method S1 section.

### Immunoblot and quantitative analyses

Immunoblot was performed according to previously described (Bedi et al., 2016, Sengupta et al., 2020). Detailed description of immunoblot technique and subsequent quantitative analyses are given in Methods S1.

### Cell free degradation assay

Two-week old WT, *abi3*, *ABI3OX-1* and *jaz1* seedlings were crushed in liquid nitrogen, resuspended in degradation buffer (50 mM Tris-MES (pH 8.0), 0.5 M sucrose, 1 mM MgCl_2_, 10 mM EDTA (pH 8.0), 5 mM DTT, 4 mM PMSF) (Kong et al., 2015). Cell debris was removed by 10 min centrifugations at 17,000g at 4°C. Total plant cell extracts prepared from WT and mutant seedlings were adjusted to equal concentration with the degradation buffer. 1 µg purified recombinant protein (ABI1_myc or ARR1_His) was incubated in 500 µg total plant cell extract for each reaction with addition of 1 mM ATP, and incubated at 25°C for different time (0, 1, 2 and 3 hour). 6x SDS loading buffer was added to stop reactions. Immunoblot analyses using anti-myc (Abcam ab32) or anti-His (BioBharati, Kolkata, India) was done to detect ABI1_myc or ARR1_His protein levels respectively. Anti-H3 immunoblot served as the loading control. For quantitative analysis, band intensity of ABI1_myc or ARR1_His were quantified using ImageJ. The starting point (0 h) was set to 1, and other points compared with it. Degradation rate was graphically plotted using the formula [Degradation = band intensity (0 hour) – band intensity (timepoint)].

## Results

### ABI3 regulates primary root length by affecting cell length in the root elongation zone

The phytohormone ABA is known to regulate primary root growth in Arabidopsis, under varying environmental conditions (Harris, 2015). ABI3 is an important transcription factor that is known to be involved in ABA signalling (Khandelwal et al., 2010, Brady et al., 2003, Finkelstein, 1994). To understand the role of ABI3 mediated ABA signalling in regulating post-embryonic primary root growth in Arabidopsis, we worked with ABI3 deletion and overexpression mutants. ABI3 deletion mutant *abi3-6*, described previously (Bedi et al., 2016, Sengupta and Nag Chaudhuri, 2020), was used for the present work and is referred to as *abi3* hereafter. Two ABI3 overexpression lines *ABI3OX-1* and *ABI3OX-2* were generated through floral dip method, as described in the Materials and Methods section. Significantly higher levels of *ABI3* expression in the roots of overexpression lines, compared to wild type was confirmed through qRT-PCR (Figure S1C). When primary root growth was observed in *abi3*, *ABI3OX-1* and *ABI3OX-2* at different days post-germination (dpg), it was observed that while absence of ABI3 (*abi3*) resulted in shorter primary root length, overexpression of ABI3 (*ABI3OX-1*, *ABI3OX-2*) caused increase in primary root length, compared to wild type (Figure 1, Figure S1A, S1B). The differences in root length became more evident with increase in the number of days post germination (Figure 1B). As both the ABI3 overexpression lines showed similar root morphology compared to wild type, further experiments hereafter was done with *ABI3OX-1* seedlings only (Figure S1A, S1B).

**Figure 1:**
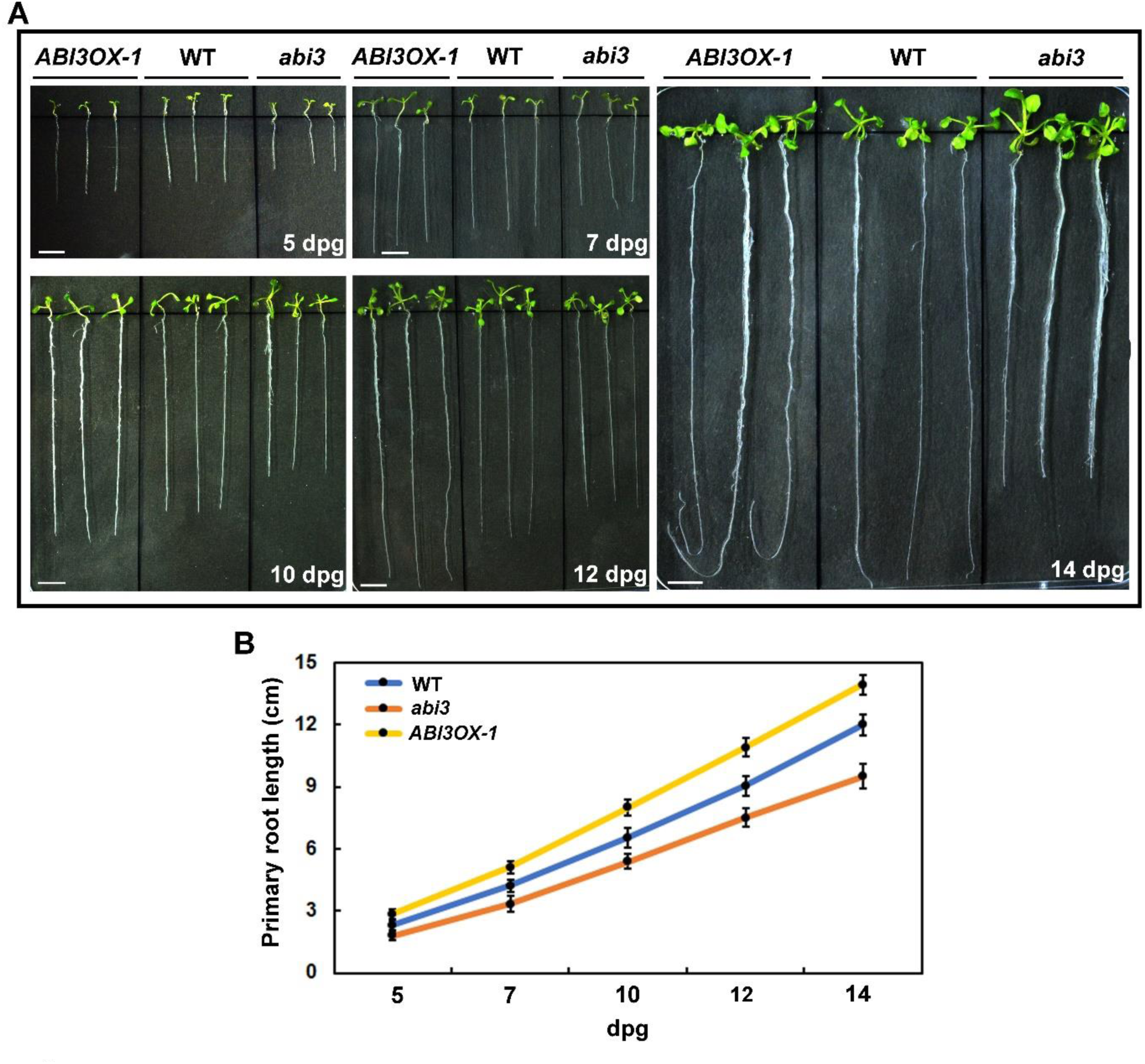
Root growth phenotype of *abi3* and *ABI3OX-1* seedlings compared to wild-type (WT). **A.** Primary root growth of wild-type (WT), *abi3* and *ABI3OX-1* grown vertically on Murashige and Skoog (MS) medium for 5, 7, 10, 12 and 14 days post germination (dpg). Scale bar: 1 cm. **B.** Primary root length measurement of WT, *abi3* and *ABI3OX-1* over time. Data represent the mean of three replicate experiments with n ≥ 20 seedlings for each replicate of each genotype with standard error of mean (SE).

Primary root is known to comprise of four distinct zones namely, meristematic zone (MZ), transition zone (TZ), elongation zone (EZ) and differentiation zone (DZ) (Verbelen et al., 2006). To understand which zone of the primary root was affected by ABI3, we did confocal microscopy of the ABI3 mutants and compared it to the wild type. As shown in Figure 2A and 2B, it was observed that in *abi3*, EZ length of the primary root was significantly reduced compared to wild type, while in *ABI3OX-1*, the EZ length was distinctly increased. Additionally, measurement of individual epidermal cell length indicated that in the *abi3* mutant, epidermal cells of EZ were significantly shorter in length, while in *ABI3OX-1*, EZ cells were significantly longer, compared to wild type (Figure 2C & 2D). The above observations indicate that absence of ABI3 affects primary root length through decrease of cell length in the root elongation zone.

**Figure 2:**
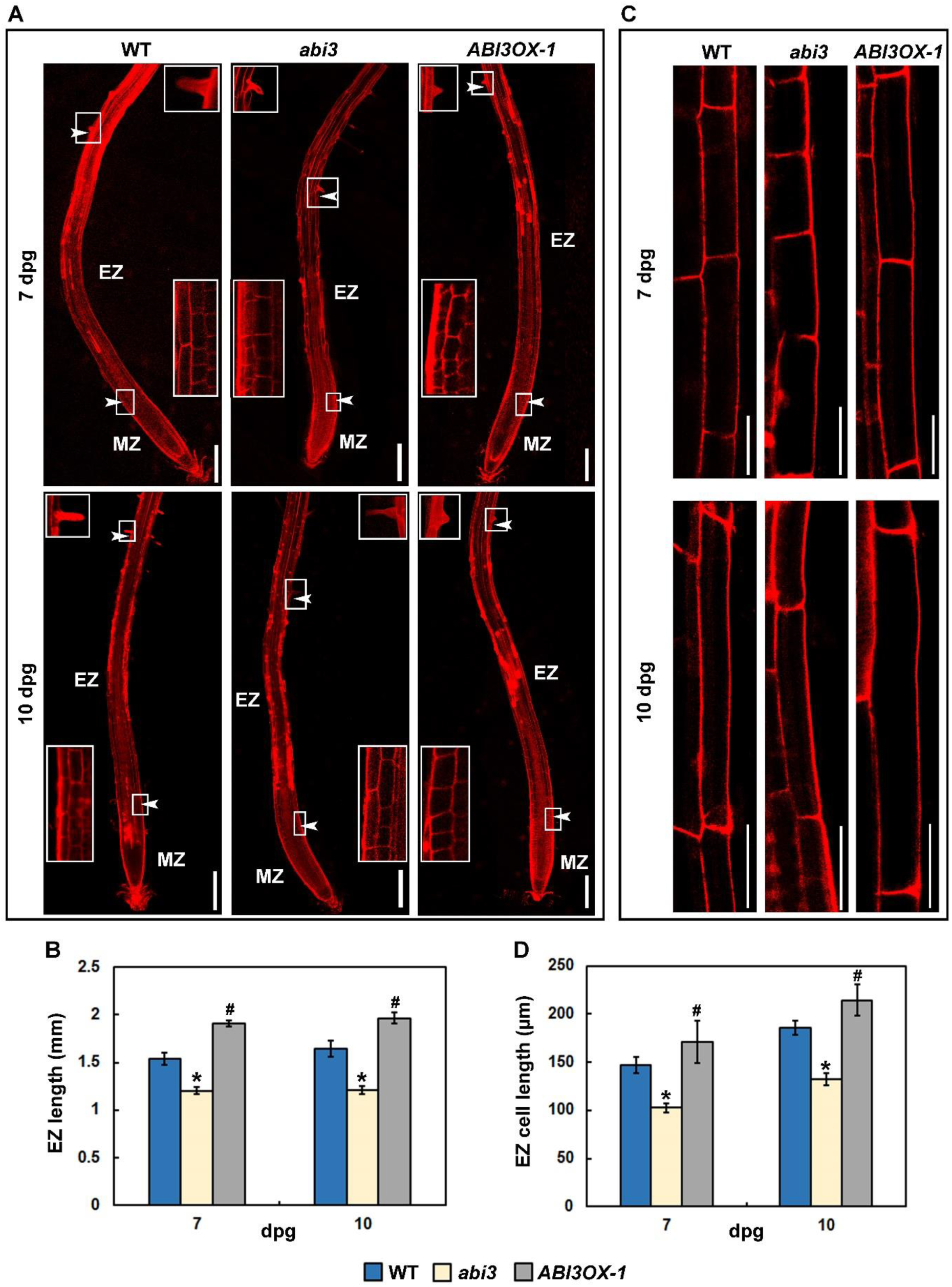
Microscopic analysis of primary root of WT, *abi3* and *ABI3OX-1.* **A.** Longitudinal view of propidium iodide stained 7- and 10-day-old WT, *abi3* and *ABI3OX-1* primary root, where the elongation zone (EZ) is represented in between the two arrowheads. The initiation of first elongated root hair, representing the differentiation zone, and the junction of the EZ and transition zone is shown in the inset of each primary root. MZ represents meristematic zone. Scale bar: 200 µm. **B.** Measurement of elongation zone length represented in (A) from 7- and 10-day-old WT, *abi3* and *ABI3OX-1* seedlings. Data represent means from three independent replicates (n ≥ 20) and error bars represent SE. Anova two-factor with replication method was used to measure the variance, and the results with *P ≤ 0.05* considered to be significant. Variance between WT and *abi3* is denoted by *, while variance between WT and *ABI3OX-1* is denoted by #. **C.** Confocal images of propidium iodide-stained epidermal root cells of the elongation zone (EZ) from 7- and 10-day-old WT, *abi3* and *ABI3OX-1* seedlings. Scale bar: 50 µm. **D.** Measurement of epidermal root cells of elongation zone represented in (C) from 7- and 10-day-old WT, *abi3* and *ABI3OX-1* seedlings. Data represent means from three independent replicates (n ≥ 20) and error bars represent SE. Anova two-factor with replication method was used to measure the variance, and the results with *P ≤ 0.05* considered to be significant. Variance between WT and *abi3* is denoted by *, while variance between WT and *ABI3OX-1* is denoted by #.

### ABI3 affects expression of JAZ genes in root

High throughput transcriptome analyses from roots of wild type and *abi3* seedlings 14 dpg, revealed differential expression of several JAZ genes, along with *MYC2* and JA biosynthesis gene *LOX2* (Figure S2, Sequence Read Archive (SRA) accession number PRJNA869323). To complement this data and delve into the molecular mechanism of ABI3 mediated effect on post-embryonic primary root growth, we isolated RNA from roots of wild type, *abi3* and *ABI3OX-1*, at different days post germination (dpg) and subjected it to qRT-PCR. It was observed that several JAZ genes such as *JAZ1*, *JAZ6*, *JAZ7*, *JAZ8* and *JAZ9* showed differential regulation in *abi3* and *ABI3OX-1*, compared to wild type (Figure 3). Of these genes, *JAZ1* consistently showed significantly higher expression in *abi3* from 5-14 dpg, compared to wild type. In *ABI3OX-1* on the other hand, *JAZ1* expression was significantly downregulated, compared to wild type. Notably, *ABI3* expression in wild type was also found to be differential at different days post germination, and got significantly downregulated during 7-14 dpg, compared to 5 dpg.

**Figure 3:**
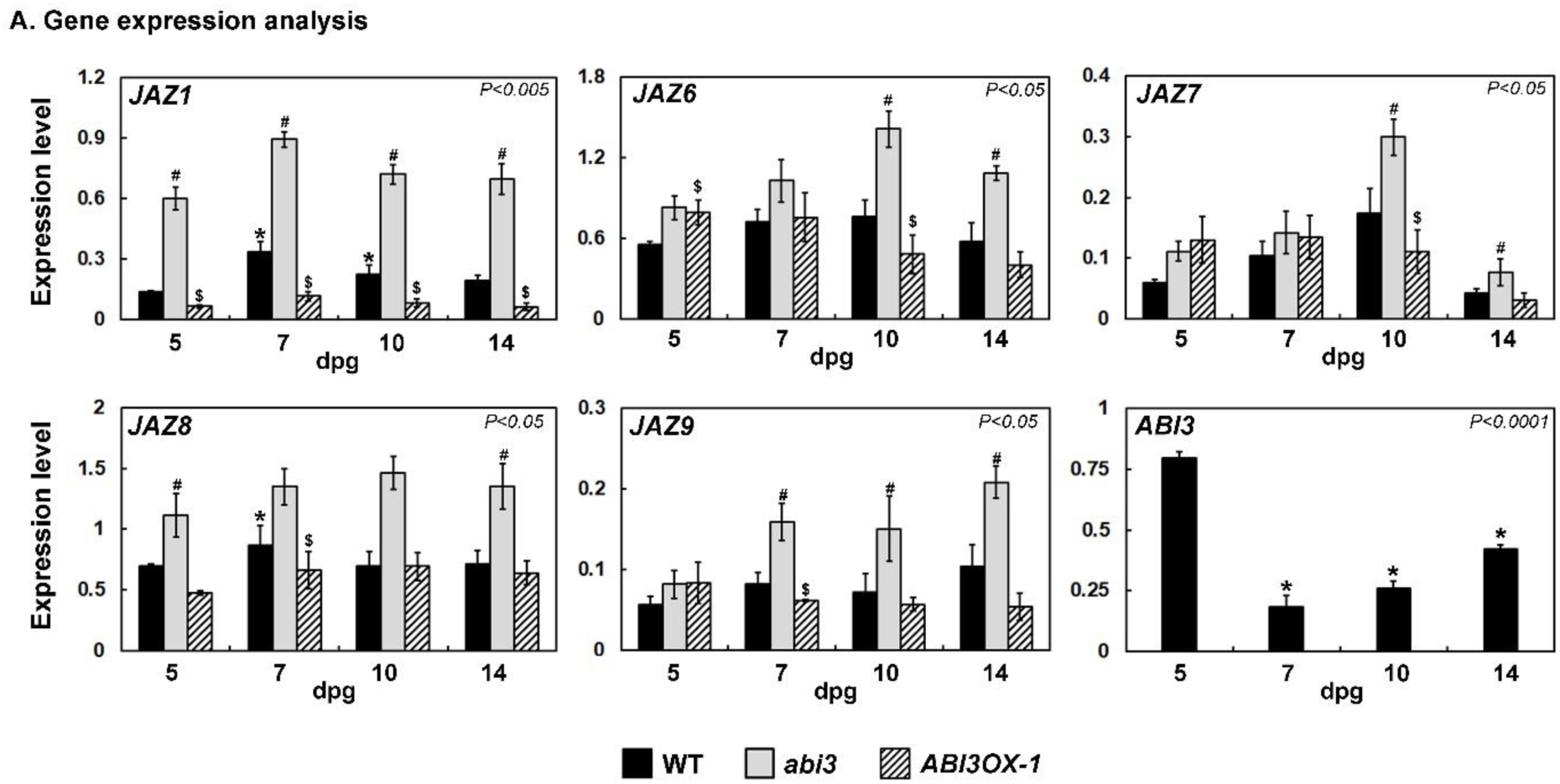
Gene expression analysis. **A.** Total RNA isolated from roots of WT, *abi3* and *ABI3OX-1* mutant seedlings at the indicated number of days post germination (dpg) and subjected to quantitative reverse transcriptase-polymerase chain reaction (qRT-PCR). All qPCRs were performed in triplicate and error bars represent SE. Anova two-factor with replication method was used to calculate the variance. Variance in between WT and *abi3* is denoted by # and in between WT and *ABI3OX-1* is denoted by $ for individual timepoint. Variance within different timepoints of WT is denoted by * on corresponding graphs. *P* value obtained is mentioned in the individual graphs.

To further understand the pattern of *JAZ1* expression in the primary root, we generated *proJAZ1::GFP* transgenic lines in wild type and *abi3* background and checked GFP expression in the primary roots, at 7dpg. As shown in Figure 4A, it was evident that *JAZ1* expression occurs in the primary root elongation zone and meristematic zone. It was interesting to note that, in consonance with the qRT-PCR data, the level of *proJAZ1::GFP* expression was higher in the primary root of *abi3,* compared to wild type. Additionally, we also generated *proABI3::GFP* transgenic line in wild type background and observed that there was distinct expression of *ABI3* in the elongation zone and meristematic zone of the primary root, at 5 and 7 dpg (Figure 4B). Similar to transcription data, *ABI3* expression in the primary root was found to be reduced at 7 dpg, compared to 5 dpg. From these observations it can be concluded that both *ABI3* and *JAZ1* expressions occur in the primary root elongation and meristematic zones, and in absence of ABI3, *JAZ1* expression is enhanced.

**Figure 4:**
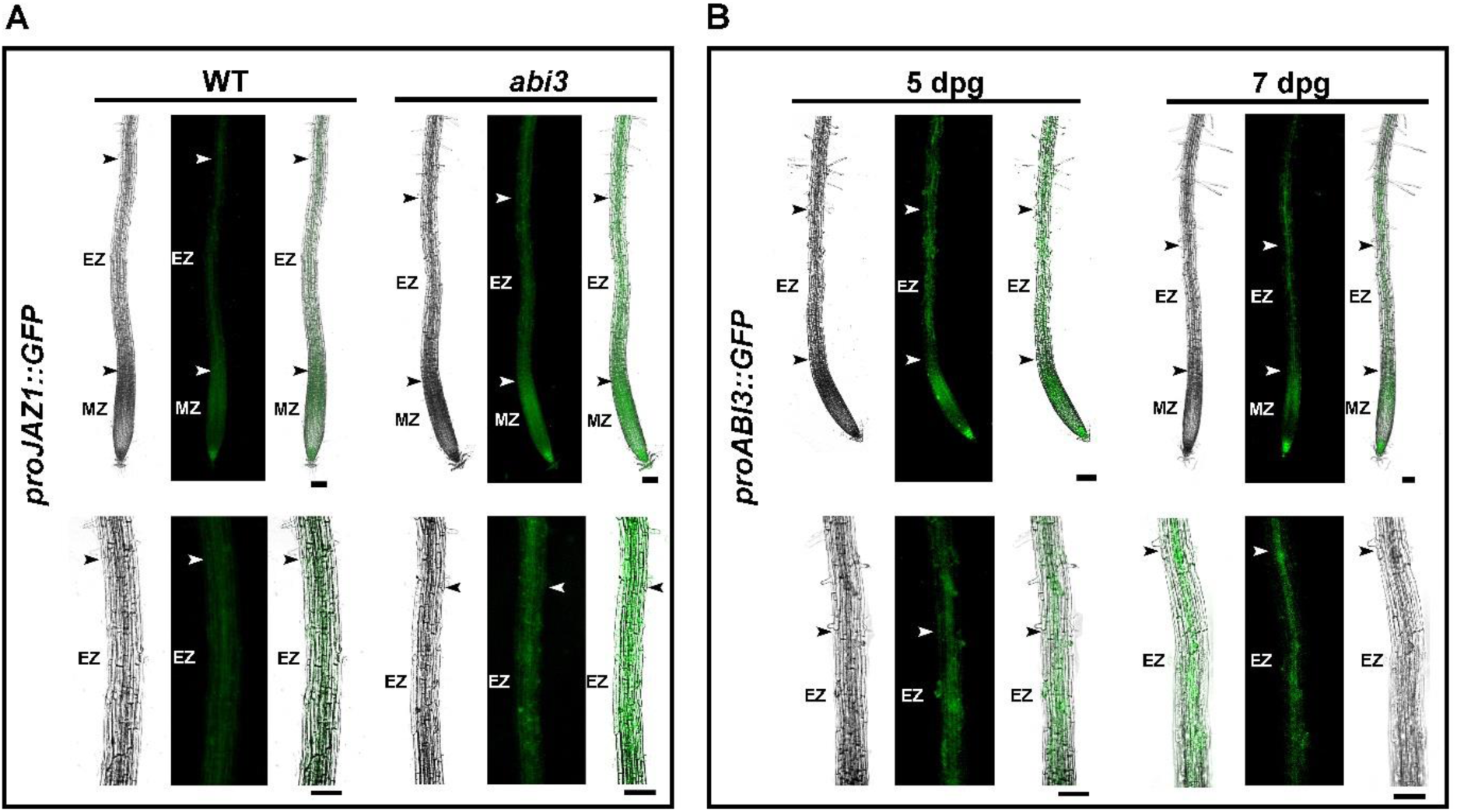
Localization of ABI3 and JAZ1 in primary root. **A.** proJAZ1::GFP expression in the primary roots of stable transgenic lines of WT and *abi3* (top panel), and in the primary root elongation zone (bottom panel), at 7 days post germination (dpg). Scale bar: 100 µm. **B.** proABI3::GFP expression in the primary root of stable transgenic line of WT (top panel), and in the primary root elongation zone (bottom panel), at 5 and 7 days post germination (dpg). Scale bar: 100 µm.

### ABI3 negatively regulates JAZ1 expression

From the above results it became apparent that *JAZ1* expression was elevated in absence of ABI3. *In silico* analyses indicated that in the *JAZ1* locus, within 500 bp upstream of the TSS there are several ABI3 binding cis-elements including ABRE (ACGTG), Sph/RY element (CATGCA) and AuxRE (TGTCTC), along with putative TATA boxes in the near vicinity (Figure 5A). Hereafter, we did ChIP analyses with wild type and *ABI3OX-1* roots 7 and 10 dpg, to understand whether ABI3 gets recruited to the promoter region of *JAZ1*. Results indicated ABI3 occupancy in the promoter region of *JAZ1*, both in wild type and *ABI3OX-1* roots. As expected, the level of ABI3 occupancy in the *JAZ1* promoter was higher in *ABI3OX-1* roots compared to wild type (Figure 5B). To correlate this data with transcription activity in *JAZ1* locus, we checked RNAPII occupancy, along with H3K4me3 and H3K27me3 histone modification marks. It is known that H3K4me3 is a mark of active transcription, while presence of H3K27me3 indicates transcription repression. We found that RNAPII occupancy in the promoter region of *JAZ1* was reduced in *ABI3OX-1* roots, compared to wild type. In consonance, H3K4me3 mark was reduced while H3K27me3 mark was enriched in the *JAZ1* promoter of *ABI3OX-1*, compared to wild type (Figure 5B). These results indicate that ABI3 occupancy in the promoter region of *JAZ1* leads to reduction in RNAPII occupancy and associated active histone modification mark like H3K4me3 that is required for active transcription. Such inhibition of ABI3-mediated *JAZ1* expression is further ensured by H3K27me3 enrichment in the upstream regulatory region, which is known to promote transcription repression.

**Figure 5:**
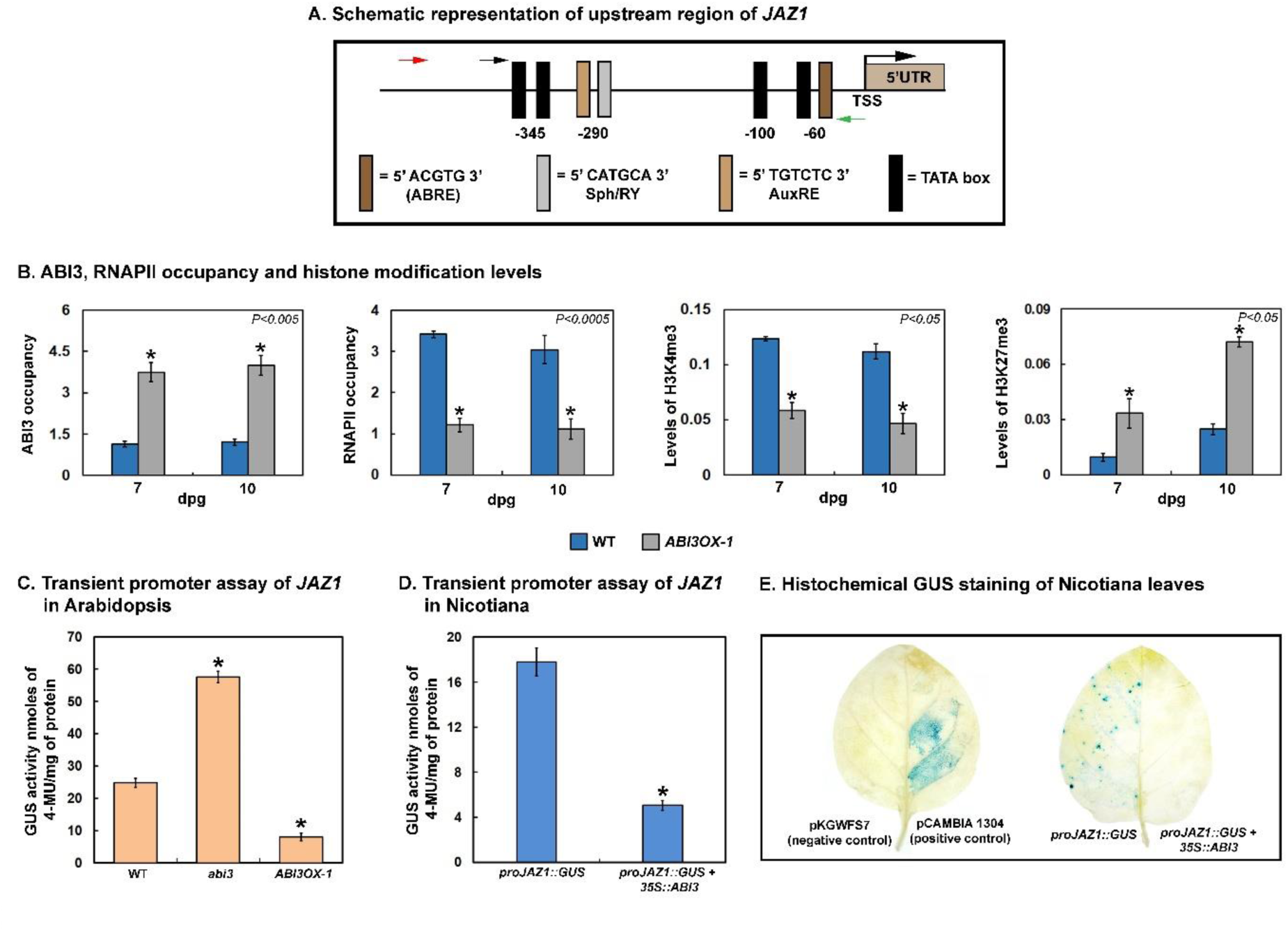
ABI3 regulates *JAZ1* promoter activity. **A.** Schematic map of *JAZ1* promoter region showing ABI3 binding sites and putative TATA boxes. The presence of the respective conserved core elements is shown relative to transcription start site (TSS). Forward primers used for ChIP-qPCR and cloning are shown as black and red arrowheads, respectively, and the common reverse primer is shown as green arrowhead. **B.** Chromatin immunoprecipitation (ChIP) assay using anti-ABI3, anti-RNAPII, anti-H3K4me3 and anti-H3K27me3 antibodies showing ABI3 and RNAPII occupancy and histone modification levels relative to H3 at promoter region of *JAZ1* gene in 7- and 10-day-old root tissue of WT and *ABI3OX-1.* The represented data are means from three independent replicates and error bars represent SE. Statistical significance tested using paired two tailed Student’s t-test. *P* value obtained is mentioned in the individual graphs and are marked by *. **C.** Leaves of 6-week-old WT, *abi3* and *ABI3OX-1 Arabidopsis thaliana* seedlings infiltrated with either resuspension buffer or *Agrobacterium tumefaciens* containing *proJAZ1::GUS*, and allowed to recover for 48 h. The infiltrated leaves were excised and subsequently GUS assay was performed. Values were normalized against resuspension buffer control. The graph represents GUS activity of *JAZ1* promoter in WT, *abi3* and *ABI3OX-1* expressed as nanomoles of 4-MU per mg of total protein. **D.** Leaves of 6-week-old *Nicotiana benthamiana* plants were either infiltrated with *proJAZ1::GUS* or co-infiltrated with *proJAZ1::GUS* and *35S::ABI3,* and after 48 h recovery infiltrated leaves were excised and used for GUS assay. The graph represents GUS activity expressed as nanomoles of 4-MU per mg of total protein. The represented data in (C) and (D) were the mean of three biological replicates over three separate occasions (n ≥ 20 leaves) with standard error bars. Student’s t-test with paired two-tailed distribution was used for statistical analysis, and *P ≤ 0.001* is denoted by *. **E.** Histochemical analysis of GUS expression in agro-infiltrated tobacco leaves. pCAMBIA 1304 and pKGWFS7 were used as positive and negative control respectively. *proJAZ1::GUS* with or without *35S::ABI3* was co-infiltrated and subsequently staining done using X-gluc.

To further understand whether ABI3 can repress *JAZ1* promoter activity, we cloned ∼700 bp fragment of *JAZ1* upstream sequences to GUS reporter gene for generating the proJAZ1::GUS construct. GUS assay was done in wild type, *abi3* and *ABI3OX-1* Arabidopsis plants, as described previously (Mandal et al., 2023, Sengupta et al., 2020). It was observed that in *abi3* mutant where functional ABI3 is absent, GUS expression increased significantly, compared to wild type. On the other hand, in *ABI3OX-1*, where ABI3 protein is overexpressed, GUS expression was significantly reduced, compared to wild type (Figure 5C). These results clearly indicate that ABI3 is a negative regulator of *JAZ1* promoter activity. This data was further complemented by *JAZ1* transactivation assay in Nicotiana, as well. For this, ABI3 cDNA was cloned under 35S promoter and the resultant construct was co-infiltrated into *Nicotiana benthamiana* leaves, along with proJAZ1::GUS construct. Results indicated that GUS activity was significantly reduced when proJAZ1::GUS was co-infiltrated with 35S::AB13 construct, compared to when proJAZ1::GUS construct was infiltrated alone (Figure 5D). This result implied that, in presence of ABI3 protein *JAZ1* promoter activity is repressed. Furthermore, histochemical GUS assay indicated that proJAZ1::GUS construct infiltrated alone in Nicotiana leaves was stained blue by X-Gluc solution, but when it was co-infiltrated along with 35S::AB13 no blue stain was observed, confirming that ABI3 supress JAZ1 promoter activity. (Figure 5E). For this assay, empty pCAMBIA1304 vector with GUS under 35S promoter was used as the positive control, while promoter-less pKGWFS7 was used as the negative control. Taken together, from the above results it can be concluded that ABI3 acts as a transcription repressor for *JAZ1* gene.

To further correlate this to root growth, we checked primary root elongation in *jaz1* deletion mutant and compared it to wild type and *ABI3OX-1.* Similar to *ABI3OX-1*, the primary root of *jaz1* mutant was found to be longer compared to wild type (Figure S3A & S3B). Confocal microscopy to check EZ of the *jaz1* primary root and the corresponding epidermal cell length indicated that the EZ size and cell length were both significantly higher in *jaz1* mutant, compared to wild type (Figure S3C-3F). From all the above results it can be therefore concluded that ABI3 mediated regulation of *JAZ1* expression controls primary root elongation. Increased expression of *JAZ1* inhibits primary root elongation, as seen in *abi3* seedlings, while reduced or absence of *JAZ1* expression as found in *ABI3OX-1* and *jaz1* seedlings respectively, lead to longer primary root and increased cell length in the EZ.

### Absence of ABI3 and presence of JAZ1 stabilizes ABI1 protein

In trying to understand how JAZ1 crosstalks with ABA signalling during post-embryonic root growth in Arabidopsis, we came across previous findings that JAZ1 interacts with ABI1, the protein phosphatase 2C, that acts as an important regulator of ABA signalling (Sun et al., 2020, Luo et al., 2020). To further correlate, we next checked ABI1 protein levels in wild type and *abi3* roots. For this, total protein was isolated from wild type and *abi3* roots with or without ABA treatment and subjected to western blot analyses using anti-ABI1 antibody. Previous studies have shown that ABI1 protein can be detected only in presence of high concentration of ABA (Kong et al., 2015). As apparent from our results, while no ABI1 protein could be detected in absence of ABA treatment, in presence of 50 μM ABA distinctly visible protein bands could be detected both in wild type and *abi3* roots (Figure 6A). It was further observed that in *abi3* roots the level of ABI1 protein was significantly higher, compared to wild type (Figure 6B). As protein level was distinctly higher in *abi3* compared to wild type, hereafter, we checked the stability of ABI1 protein in presence and absence of functional ABI3, through cell-free degradation assay. For this, recombinant myc-tagged ABI1 protein was expressed in *E.coli* and subsequently purified. The recombinant protein was incubated for various time periods with total plant cell extracts isolated from wild type and *abi3* mutant seedlings and the resultant western blot was probed with anti-myc antibody. H3 protein was used as the normaliser. Results from the cell-free degradation assay indicated that, incubation of the ABI1 protein with *abi3* plant cell extract led to a much slower rate of degradation of the protein, in comparison to wild type (Figure 6C & 6D). This implied that in absence of ABI3, ABI1 protein remains more stabilized. To further correlate, we checked ABI1 protein stability in presence of *ABI3OX-1* and *jaz1* plant cell extracts, as well. It was found that overexpression of ABI3 or absence of JAZ1, led to much higher rate of degradation of ABI1 protein, compared to *abi3*. In fact, the degradation rate of ABI1 in presence of *ABI3OX-1* or *jaz1* plant cell extracts was even higher than that of wild type (Figure 6C & 6D). These observations implied that presence of ABI3 or absence of JAZ1, causes reduced stability of ABI1 protein, while absence of functional ABI3 increases it. As a control, cell-free degradation assay was done with recombinant ARR1 protein using wild type, *abi3*, *ABI3OX-1* and *jaz1* plant cell extracts. It was found that degradation rate of ARR1, was similar in presence of all the plant cell extracts used (Figure 6C & 6E). To sum up, it can be concluded that while absence of ABI3 leads to higher *JAZ1* expression, overexpression of ABI3 or absence of JAZ1, leads to reduced stability of ABI1 protein. Therefore, in *abi3* plants where *JAZ1* expression levels are high, ABI1 protein is more stable. It is possible that in *abi3* where higher expression of *JAZ1* occurs, interaction of JAZ1 with ABI1, prevents degradation of the latter.

**Figure 6:**
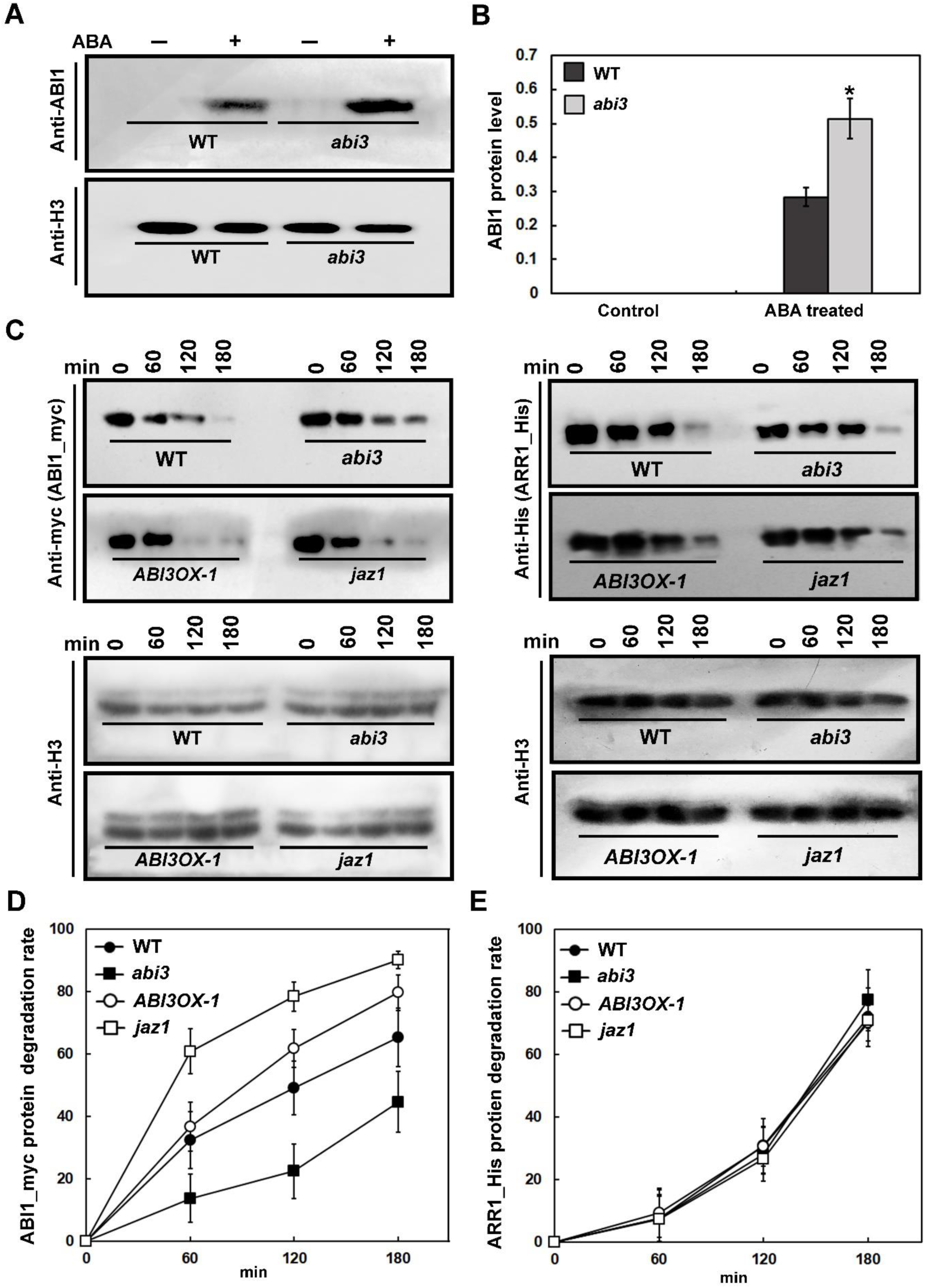
ABI1 protein stability in absence of ABI3. **A.** 7-day-old WT and *abi3* seedlings were treated with or without 50 µM ABA for 6 h, the root samples were harvested and total proteins was isolated. The proteins were subjected to immunoblotting analysis with anti-ABI1 antibody (top panel). H3 protein was used as a loading control (bottom panel). **B.** Quantitative analyses of the band intensities in (A). Band intensities of ABI1 was normalized with respect to H3 of each sample and were graphically plotted. The represented data are means from three independent replicates and error bars represent SE. Student’s t-test with paired two-tailed distribution was used for statistical analysis, and *P ≤ 0.05* between WT (+ABA) and *abi3* (+ABA) is denoted by *. **C.** Cell-free degradation assay showing the degradation rates of ABI1_myc (top left panel) and ARR1_His (top right panel) incubated with the total plant cell extract of WT, *abi3*, *ABI3OX-1* and *jaz1*, respectively. The degradation rate of ABI1_myc and ARR1_His was detected by anti-myc and anti-His antibodies respectively. H3 protein was used as loading control (bottom panel). **D.** Quantitative analyses of the degradation rate of ABI1_myc protein. Band intensities from each time point was subtracted from 0 min of each assay for different mutant lines. Error bars indicate the SE (n=3 biological replicate). **E.** Quantitative analyses of the degradation rate of ARR1_His protein. Band intensities from each time point was subtracted from 0 min of each assay for different mutant lines. Error bars indicate the SE (n = 3 biological replicate).

### ABI3 mediated regulation of primary root length involves ABI1 function

To understand the role of ABI1 in post-embryonic primary root growth, we checked the root morphology of ABI1 deletion (*abi1*) mutant seedlings 7 and 14 dpg. As shown in Figure S4, we found that *abi1* seedlings had longer primary root, compared to wild type, similar to *ABI3OX-1* and *jaz1* seedlings. Confocal microscopy revealed that the EZ length and its individual epidermal cell length is higher in the primary root of *abi1* mutant, compared to wild type (Figure S4C-4F). From these observations it is evident that, in absence of ABI1, cell length in the EZ of primary root is increased. Length control of cells, especially in the roots is a function of auxin transport mediated apoplastic pH change and integrally involves the action of plasma membrane H^+^ATPase family of proteins (Gámez-Arjona et al., 2022, Du et al., 2020). It is known that auxin-dependent function of the plasma membrane H^+^ ATPases leads to acidification of the apoplast that in turn activates cell wall loosening enzymes and contributes to cellular elongation (Hager, 2003, Samalova et al., 2023, Cosgrove, 1999). Previous studies have shown that ABI1 can regulate phosphorylation status of plasma membrane H^+^ATPase AHA2 (Miao et al., 2021). While phosphorylated AHA2 is functionally active, dephosphorylation of it by ABI1 leads to its functional inactivation (Duby and Boutry, 2009, Sondergaard et al., 2004). To further establish the relation between ABI1 and AHA2 we isolated protein from the roots of *abi3* and wild type seedlings, 7 and 10 dpg. We first performed a western blot analysis using antibody against plasma membrane H^+^ ATPase and found that the protein expression level was similar in roots of wild type and *abi3* (Figure 7A & 7B). AHA2 is known to get phosphorylated at its penultimate residue Thr^947^ located at its C-terminal end (Duby and Boutry, 2009, Sondergaard et al., 2004). Therefore, to specifically understand the status of AHA2 phosphorylation in wild type and *abi3* we next performed western blot analysis using a peptide antibody against phosphorylated Thr^947^. As shown in Figure 7A and 7B, the level of AHA2 phosphorylation was significantly reduced in *abi3*, compared to wild type, both at 7 and 10 dpg. From the above results it is therefore evident that in absence of ABI3, AHA2 is predominantly in a dephosphorylated state. Such a state of reduced phosphorylation may inhibit protonation of the apoplast, thus leading to inhibition of root cell elongation.

**Figure 7:**
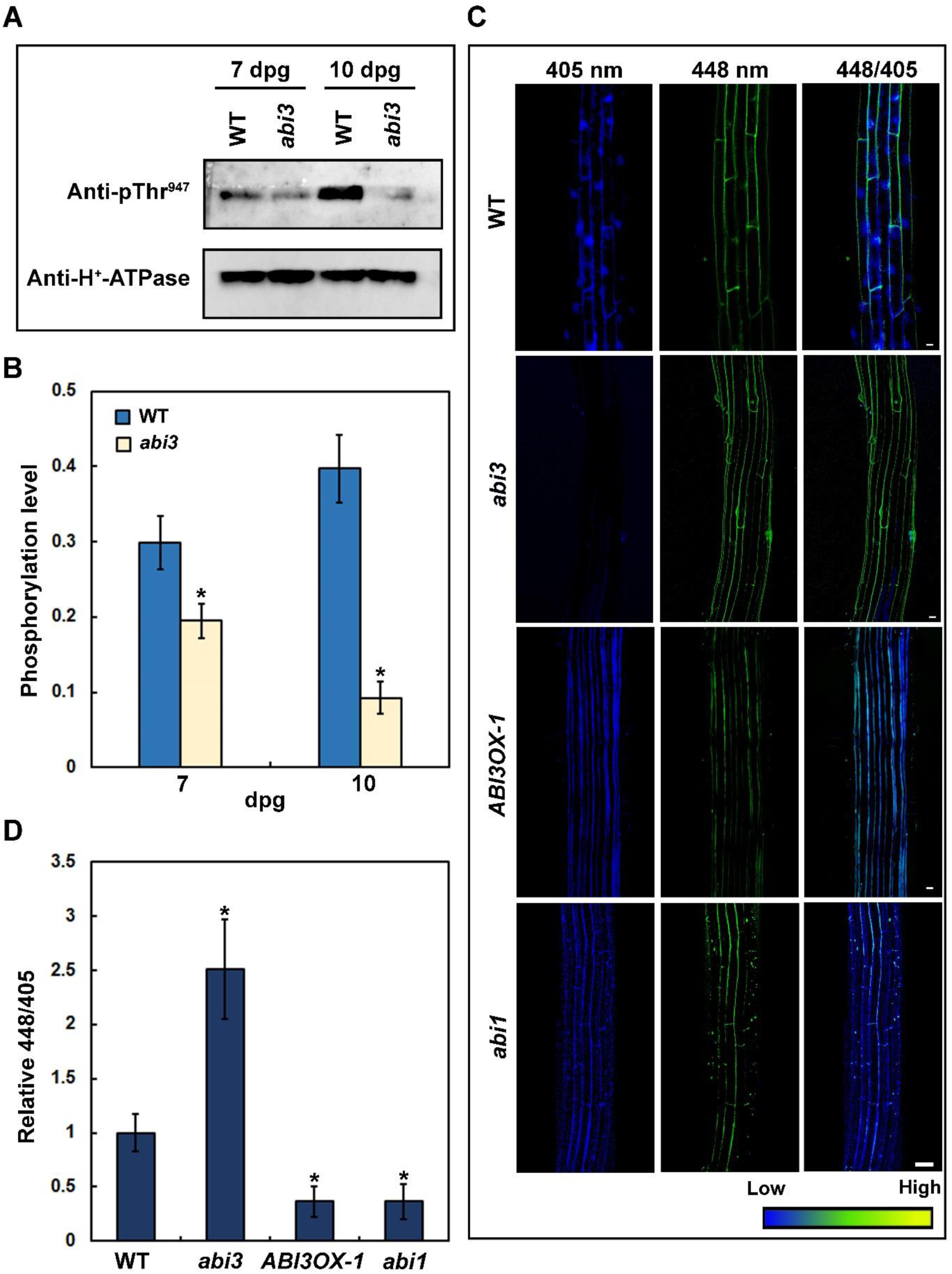
ABI3 regulates apoplastic pH of the elongation zone. **A.** Western blot detection of phosphorylated H^+^-ATPase in the root samples of 7- and 10-day-old WT and *abi3* seedlings (top panel). AHA protein levels were determined using anti-H^+^-ATPase antibody (bottom panel). **B.** Quantification of the phosphorylation level of the H^+^-ATPase represented in (A). The phosphorylation level of AHA2 was quantified as the ratio of the signal intensity from phosphorylated AHA2 to that of H^+^-ATPase. Values are means ± SE, n =3 independent experiments. Student’s t-test with paired two-tailed distribution was used for statistical analysis, and *P ≤ 0.05* is denoted by *. **(C)** Comparison of the apoplastic pH in the primary root elongation zone of 5-day-old WT, *abi3*, *ABI3OX-1* and *abi1* mutants, visualized by HPTS staining. The fluorescent signal represents protonated HPTS (405) (Excitation 405 nm, emission peak 514 nm), deprotonated HPTS form (448) (Excitation 448 nm, emission peak 514 nm) and the merge of 448/405. Color code (blue to yellow) depicts (low to high) 448/405 intensity. Scale bar: 50 µm. **(D)** Quantification of fluorescent ratio of deprotonated HPTS to protonated HPTS in the epidermal cells of the elongation zone, as indicated in (C). The fluorescent ratio of WT was set to 1.0 and the mutant ratio was normalized to WT. Data are mean ± SE from three independent experiments (n = 15 cells from 5 roots per genotype).

8-hydroxypyrene-1,3,6-trisulfonic acid trisodium salt (HPTS) is a fluorescence pH indicator that can assess apoplastic pH of root cells (Barbez et al., 2017). Fluorescent signals from the protonated and deprotonated HPTS may be detected under specific excitation and emission wavelengths using a confocal microscope. To correlate cell length differences in the EZ of *abi3* and wild type to acidification of apoplast, we next stained *abi3*, *ABI3OX-1* and wild type roots with HPTS, using protocol as mentioned in the Materials and Methods section. As shown in Figure 7C and 7D, images were taken at 405 nm and 448 nm excitation after HPTS staining, the 448/405 ratio was analysed, and compared to wild type. Higher 448/405 value means higher apoplastic pH, while a lower value indicates increased protonation or acidification of the apoplast, compared to wild type. From our observations it was evident that compared to wild type, *ABI3OX-1* roots had lower apoplastic pH, while that of *abi3* was significantly higher. Therefore, in absence of ABI3, ABI1-mediated reduced phosphorylation of AHA2, supress acidification of apoplastic pH leading to its alkalinization. To further confirm this when roots of *abi1* mutant seedlings were HPTS stained, it was found that the 448/405 value or apoplastic pH was lower, compared to wild type. This indicated that in absence of ABI1, AHA2 mediated protonation of root apoplast occurs more efficiently (Figure 7C & 7D). Our result is similar to a previous finding that showed higher levels of phosphorylated Thr^947^ and consequent lower apoplastic pH in PP2C quadruple mutant, compared to wild type (Miao et al., 2021). Taken together, it seems pertinent to conclude that in *abi3,* where ABI1 is more stabilised, predominantly dephosphorylated form of AHA2 exists, which results in an increase in apoplastic pH leading to reduced cell elongation in the primary root. This leads to shortening of the primary root EZ and reduced primary root elongation in *abi3* seedlings, compared to wild type.

## Discussion

Each designated zone of an Arabidopsis primary root is an effective output of interplay between specific set of hormones (Liu et al., 2017, Zluhan-Martínez et al., 2021, Petricka et al., 2012). The elongation zone or EZ of the primary root is known to involve signalling of various hormones like auxin, ABA, JA along with ethylene (Chen et al., 2011, Du et al., 2020, Miao et al., 2021, Růzicka et al., 2007, Sun et al., 2018). In trying to decipher the role of ABA in determining post embryonic EZ development in primary root, we worked with ABI3, a critical transcription factor known to mediate ABA signalling and its crosstalk with other hormones (Liu and Timko, 2021, Nag et al., 2005, Pan et al., 2020). Previously we have shown that in absence of ABI3 (*abi3-6* mutant), *de novo* root organogenesis is hindered in Arabidopsis callus cells (Sengupta and Nag Chaudhuri, 2020). Here, we worked with *abi3-6*, which is known to be the most severe ABI3 deletion mutant (Nambara et al., 1994, Bedi et al., 2016, Sengupta and Nag Chaudhuri, 2020), along with ABI3 overexpression line *ABI3OX-1*. From the morphological analyses it was evident that, absence of ABI3 led to shortening of the primary root, compared to wild type, while overexpression of ABI3 caused increased primary root growth. It implied that ABI3 has a distinct role to play during post-embryonic primary root growth. Microscopic analyses revealed that ABI3 affected growth in the EZ of the primary root. More specifically, the length of epidermal cells in the EZ were shortened in absence of ABI3, and were longer in *ABI3OX-1* seedlings, when compared to wild type. We therefore argued that ABI3 directly or indirectly affects cell elongation in the EZ of Arabidopsis primary root. JA is a crucial plant hormone that is known to be involved in several developmental regulation, in crosstalk with other hormones like ABA (Liu and Timko, 2021, Pan et al., 2020). To delve into the molecular basis of ABI3 mediated control of primary root length, from our high throughput data we noted that several genes involved in JA signalling and biosynthesis were differentially regulated in absence of ABI3. Low throughput transcription analyses indicated that, *JAZ1* was most significantly differentially regulated in roots of mutant seedlings at different days post germination. *JAZ1* expression was upregulated in absence of ABI3, and significantly downregulated when ABI3 was overexpressed. Interestingly, both *ABI3* and *JAZ1* were found to get expressed in the EZ of the primary root. In wild type cells, *JAZ1* expression in the primary root was low, indicating that under control conditions, expression of *JAZ1* is tightly regulated. However, in absence of ABI3, expression of *JAZ1* in the primary root increased substantially, further endorsing that ABI3 critically regulates *JAZ1* expression during primary root development. Thus, ABI3 mediated expression regulation of *JAZ1* during EZ development, is part of a probable crosstalk between ABA and JA signalling in the process. Previously it has been reported that ABI3 interacts with JAZ1 and 35S::JAZ1-ΔJas seedlings that lack the interaction, shows strong ABA insensitive phenotype during seed germination (Pan et al., 2020). While these observations indicated an interplay between JAZ1 and ABI3, our results show for the first time that ABI3 can regulate *JAZ1* at the transcript level during primary root development.

As several ABI3 binding sites were present in the upstream regulatory region of the *JAZ1* locus, we found it pertinent to investigate whether ABI3 occupies *JAZ1* promoter region. ChIP analyses confirmed recruitment of ABI3 to the *JAZ1* promoter region and further indicated that in presence of ABI3, RNAPII occupancy was significantly impaired. This appeared plausible, as *in silico* analyses indicated that the ABI3 binding cis-elements were located very near to the putative TSS of the *JAZ1* locus. It is thus probable that ABI3 occupancy poses a steric hindrance to RNAPII recruitment at the *JAZ1* promoter region, resulting in transcription inhibition. ABI3 therefore renders *JAZ1* locus impaired for transcription. To further confirm, we checked two transcription associated histone modification marks in the *JAZ1* locus. Our results confirmed that in presence of ABI3, *JAZ1* promoter region showed enrichment of H3K27me3, which is known to be a mark of transcription repression (Kundu et al., 2017, Cai et al., 2021, Hosogane et al., 2016, Zhang et al., 2020). Additionally, presence of ABI3 correlated with reduced presence of H3K4me3 mark, which is essentially a mark of active transcription (Wozniak and Strahl, 2014, Ding et al., 2011). It is therefore evident that in presence of ABI3, chromatin milieu at the *JAZ1* locus is not conducive for active transcription. To further establish that ABI3 works as a transcription repressor of *JAZ1*, we performed transient promoter activation assay. Assay with proJAZ1::GUS construct revealed that in presence of ABI3, promoter activity of *JAZ1* was repressed, and the repression got alleviated in absence of ABI3. It thus implied that ABI3 and JAZ1 function in an antagonistic manner, with ABI3 working as a negative regulator of *JAZ1*. This was further apparent from the morphological studies, where we observed that unlike *abi3*, *jaz1* mutants show increased primary root growth. Congruently, EZ size and the length of epidermal EZ cells were higher in the *jaz1* mutant compared to wild type. Thus, while absence of ABI3 cause reduction in primary root length, absence of JAZ1 increase primary root length. It is therefore apparent, that ABI3 mediated primary root length control involves a negative regulation of JAZ1 expression.

To further correlate JAZ1 to ABA signalling we found that JAZ1 is known to interact with ABI1 protein (Pan et al., 2020, Sun et al., 2020). To connect this to our findings, when ABI1 protein level was checked in *abi3* mutant, it was found that in absence of ABI3, ABI1 protein level was higher than wild type. Cell free degradation assay indicated that in absence of ABI3, ABI1 protein was more stabilized as evident by its lower rate of degradation, compared to wild type. Importantly, overexpression of ABI3 or absence of JAZ1, caused a much faster rate of ABI1 degradation, relative to wild type. From these observations, it can be presumed that in absence of ABI3 and effectively higher JAZ1 levels, ABI1 protein is more stable. On the other hand, overexpression of ABI3 or absence of JAZ1, renders ABI1 more vulnerable to degradation. Previously, it has been shown that ABI1 is degraded upon ABA treatment, through interaction with PUB12/13, which ubiquitinates ABI1 for proteasomal degradation (Kong et al., 2015). It is possible that interaction of JAZ1 with ABI1 somehow impedes such interaction consequently leading to its stabilization and that is why we observed increased stability of ABI1 in *abi3* seedlings that express higher levels of *JAZ1*. The exact mechanism of JAZ1-mediated ABI1 stabilisation however needs further investigation and may be an interesting avenue of future research. Hereafter, it was pertinent to correlate how increased stability of ABI1 could affect EZ length during primary root growth. Previous studies have shown that ABI1 can dephosphorylate the plasma membrane H^+^ATPase AHA2, thus rendering it functionally inactive (Miao et al., 2021). Plasma membrane H^+^ATPAses function in controlling root cell length through auxin signalling. H^+^ATPase mediated proton extrusion is integrally connected to auxin transport and such acidification of the root cell apoplast is known to activate cell wall loosening enzymes like those belonging to the Expansin family (Cosgrove, 1999, Samalova et al., 2023). Such protonation of the root cell apoplast consequently regulates cell elongation and controls primary root EZ length (Gámez-Arjona et al., 2022, Staal et al., 2011). Here we found that, despite the expression level of H^+^ATPase being similar in both wild type and *abi3* roots, in *abi3* AHA2 phosphorylation levels were significantly lower, compared to wild type. This clearly indicated that in absence of ABI3, when ABI1 protein has higher stability compared to wild type, the phosphorylation level of AHA2 is distinctly reduced. This consequently affected the functional activity of AHA2. This was further evident from the HPTS staining of *abi3* and wild type roots, which gives a measurement of root cell apoplastic pH. Results clearly indicated that absence of ABI3 caused increased deprotonation, while absence of ABI1 caused increased protonation of root cell apoplast, compared to wild type. Therefore, in absence of ABI3, when ABI1 protein is more stable, reduced phosphorylation of AHA2 cause decreased protonation of the root cell apoplast. This phenomenon was further confirmed by the fact that in *ABI3OX- 1* roots the cellular apoplastic pH was significantly reduced, compared to both wild type and *abi3*. Thus, reduced protonation of cellular apoplast in *abi3* primary roots, consequently inhibits root cell elongation, resulting in shortened EZ and primary root length.

To summarize, as shown in Figure 8, ABI3 plays a critical role in regulating primary root length through regulation of two important hormone responsive factors JAZ1 and ABI1. ABI3 augments primary root length through negative regulation of *JAZ1* expression by occupying its promoter region and generating a chromatin landscape that is non-conducive for active transcription. Furthermore, absence of ABI3 stabilizes ABI1 protein, possibly through mediation of JAZ1. Increased stability of ABI1 leads to reduced phosphorylation of AHA2, the plasma membrane H^+^-ATPase. De-phosphorylated state of AHA2, compromises its functional activity leading to decreased protonation of *abi3* root cell apoplasts. Such alkalinization of the apoplast inhibited cell elongation, and decreased EZ length of the primary root causing shortening of the root length in *abi3*. ABI3 therefore impairs ABI1 function to promote AHA2- mediated protonation of root cell apoplast effectively regulating cell length. Thus, to conclude, ABI3 promotes primary root elongation by controlling cell length at the EZ. Such a finding is of significance when it comes to understanding how stress resilient plants may be generated through manipulation of root architecture and justifies why absence of ABI3, renders Arabidopsis plants hypersensitive to dehydration stress, compared to wild type (Bedi et al., 2016).

**Figure 8:**
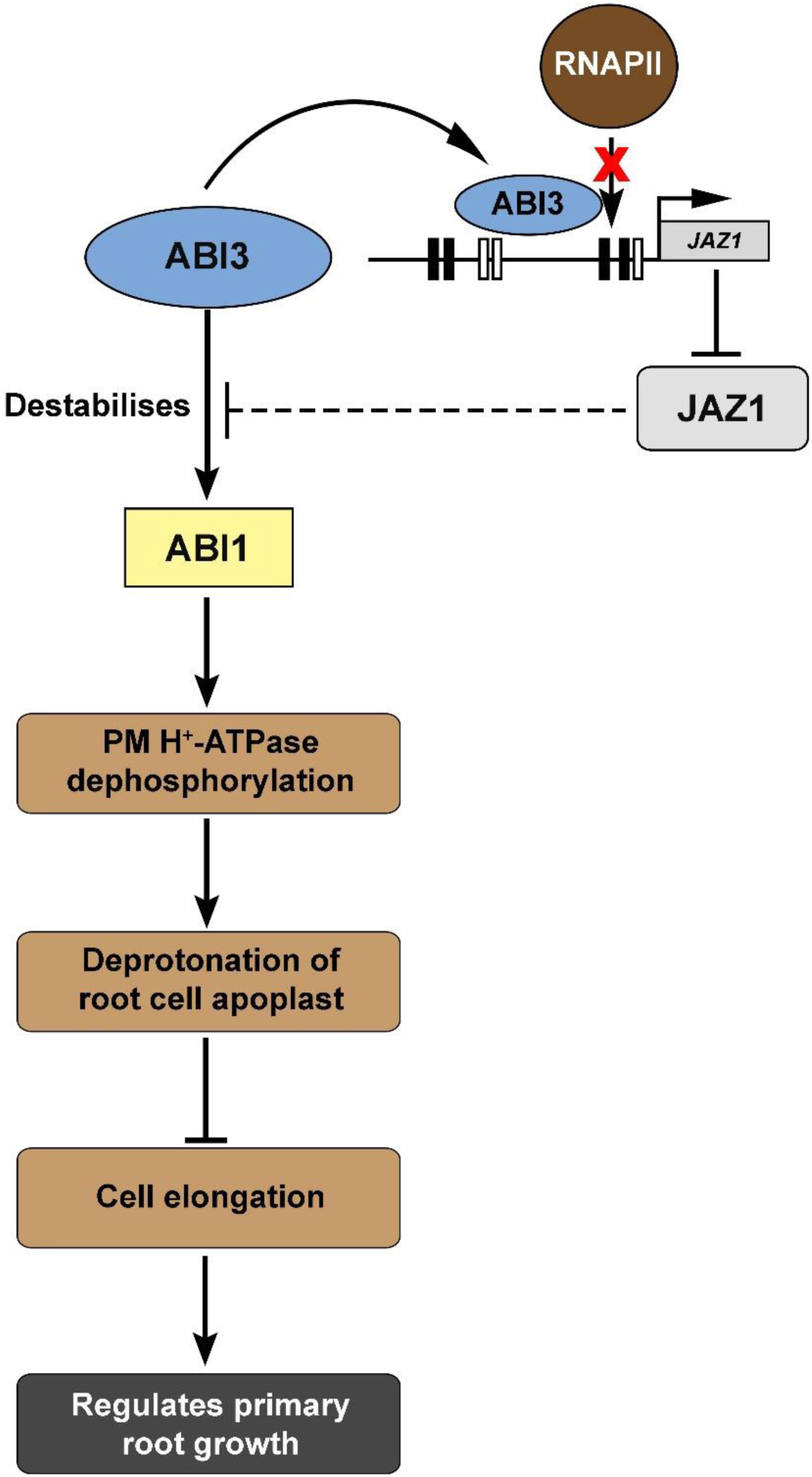
A schematic representation illustrating the function of ABI3 in regulating primary root elongation zone. ABI3 occupies *JAZ1* upstream regulatory region to repress *JAZ1* expression. ABI3 regulates ABI1 protein stability that is possibly reversely affected by JAZ1. ABI1 dephosphorylates and deactivates PM H^+^-ATPase, resulting in alkalinization of root cell apoplasts. This consequently leads to reduction in cell length within the elongation zone and negatively regulates primary root growth.

## Acknowledgement

The authors are sincerely thankful to Dr. Elke Barbez, CIBSS, Freiburg, Germany for helping with HPTS staining of Arabidopsis roots. The authors sincerely acknowledge Dr. Dibyendu Das, IISER, Kolkata for generously providing the unphosphorylated peptide of AHA2. The authors are grateful to Dr Senjuti Sinha Roy, NIPGR, India for providing with the vectors pKGWFS7 and pK7WG2D as generous gifts. The authors express their gratitude to Prof. Maitrayee Dasgupta, University of Calcutta, India, for generously providing *P19* containing plasmid and ATP (Sigma Aldrich). The authors are thankful to Prantik Saha and Confocal Microscopy facility of Bose Institute, Kolkata for assisting with confocal microscopy. This work was supported by SERB [Grant no: SPG/2021/002264] to Dr. Ronita Nag Chaudhuri; by the CSIR fellowship [08/548(0010)/2021-EMR-I] to Saptarshi Datta and by DBT [Grant no: BT/INF 22/SP41296/2020] to St. Xavier’s College, Kolkata.

## Competing Interest

The authors declare no competing interest. Contents of this manuscript are solely the responsibility of the authors and do not necessarily represent the official views of the funding agency.

## Authors Contribution

RNC, SD and DM designed the experiments; SD, DM and SM performed the experiments; RNC, SD and DM analysed the results. Manuscript written by RNC, SD, DM and SM. All the authors approve of the final version of the manuscript.

## Data Availability

All data that supports the findings presented in this article are available within the manuscript and the accompanying supplementary materials. Raw data from the RNA-seq samples have been submitted to the National Centre for Biotechnology Information (NCBI) Sequence Read Archive (SRA) database (# PRJNA869323). The data will be publicly available and accessible at https://www.ncbi.nlm.nih.gov/sra/PRJNA869323 after the indicated release date 09-12- 2024. For any further information regarding the RNA-seq data, please contact the corresponding author.

## Supporting Information

**Figure S1:** *ABI3OX* mutant lines exhibit longer primary root growth phenotype

**Figure S2:** Comparative gene expression analyses from WT and *abi3* roots 14 dpg

**Figure S3:** *jaz1* mutant line exhibit longer primary root growth phenotype

**Figure S4:** *abi1* mutant line exhibit longer primary root growth phenotype

**Figure S5:** ChIP-PCR for negative controls of ABI3 and RNAPII occupancy

**Method S1:** Detail description of the methods used in this study

**Table S1:** List of primers

## Notes

### Competing Interest Statement

The authors have declared no competing interest.

