## Supporting Information for "ABI3 regulates ABI1 function to control cell length in primary root elongation zone"

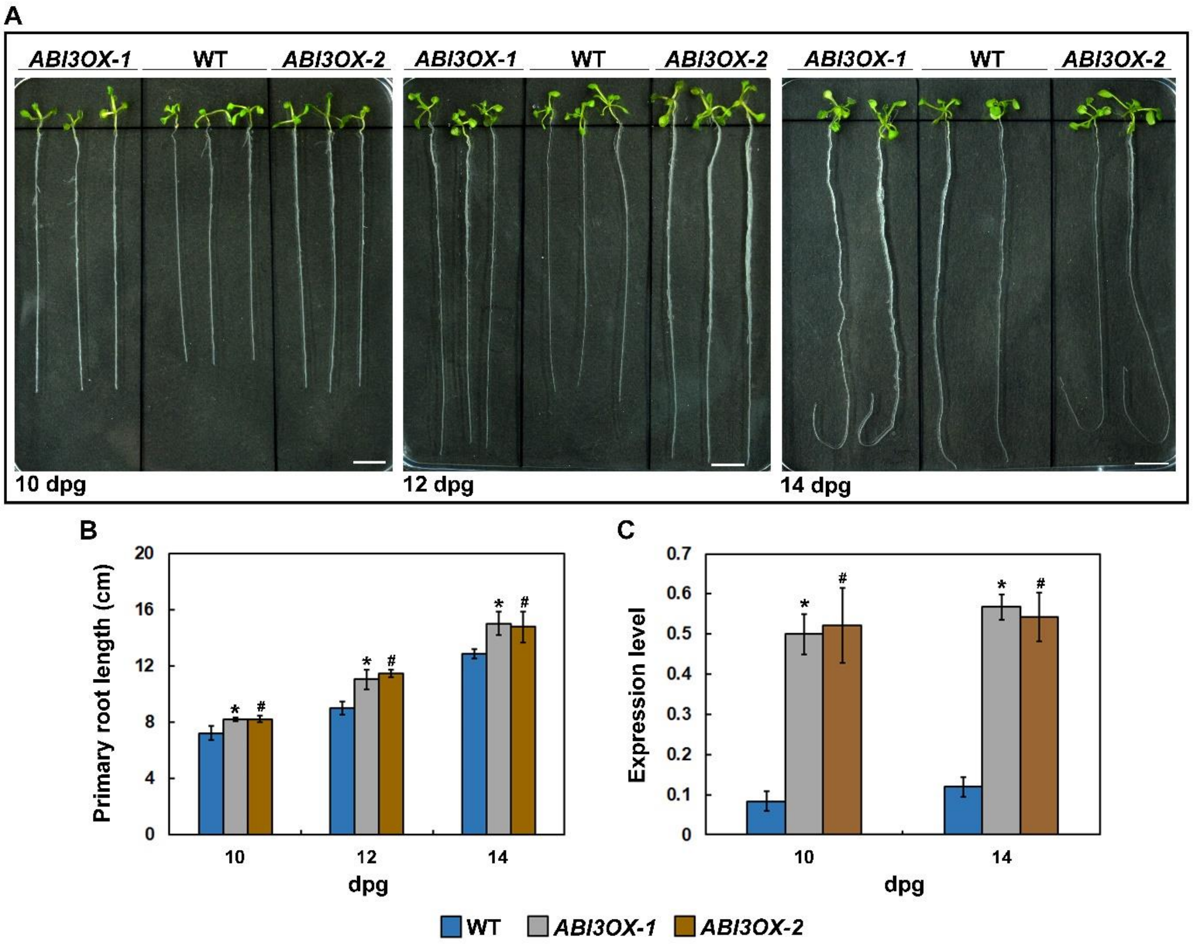


Figure S1

**Figure S1: *ABI3OX* mutant lines exhibit longer primary root growth phenotype. A.** Primary root growth of wild-type (WT), *ABI3OX-1* and *ABI3OX-2* grown vertically on Murashige and Skoog (MS) medium for 10, 12 and 14 days post germination (dpg). Scale bar: 1 cm. **B.** Primary root length measurement of WT, *ABI3OX-1* and *ABI3OX-2* over time. Data represent the mean of three replicate experiments with $n \geq 20$ seedlings for each replicate of each genotype with standard error of mean (SE). Anova two-factor with replication method was used to measure the variance, and the results with $P \leq 0.05$ considered to be significant. Variance between WT and *ABI3OX-1* is denoted by *, while variance between WT and *ABI3OX-2* is denoted by #. **C.** Expression levels of *ABI3* in roots of 10- and 14-day-old WT, *ABI3OX-1* and *ABI3OX-2* seedlings, checked by qRT-PCR using *GAPDH* as the internal control. Error bars represent SE. Anova two-factor with replication method was used to measure the variance, and results with $P \leq 0.05$ considered to be significant. Variance between WT and *ABI3OX-1* is denoted by *, while variance between WT and *ABI3OX-2* is denoted by #.

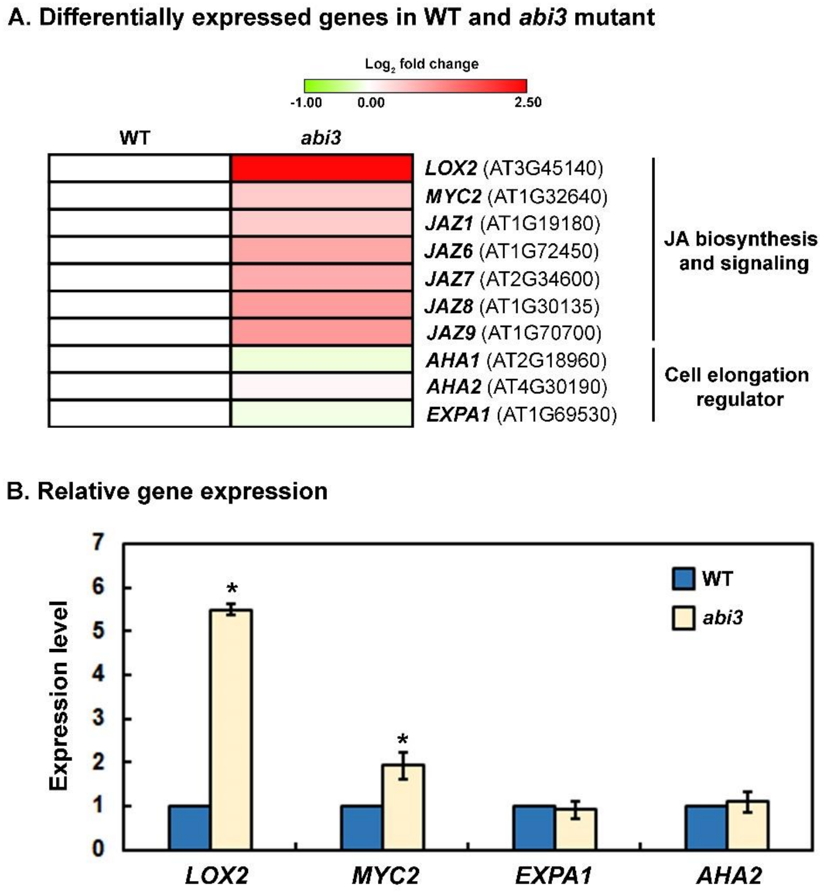


Figure S2

**Figure S2: Comparative gene expression analyses from WT and *abi3* roots 14 dpg. A.** Heat map generated (MEV 4.6.0 software) from RNA-seq analyses, representing $\log_2$ fold change of transcripts (*abi3*/WT), for genes primarily involved in jasmonic acid pathway and related to cell elongation. **B.** Validation of RNA-seq data using qRT-PCR. The data represented are mean of three biological replicates with standard error bars (SE). Statistical significance was tested using paired two-tailed student's t-test and results with *P<0.05* are marked by *.

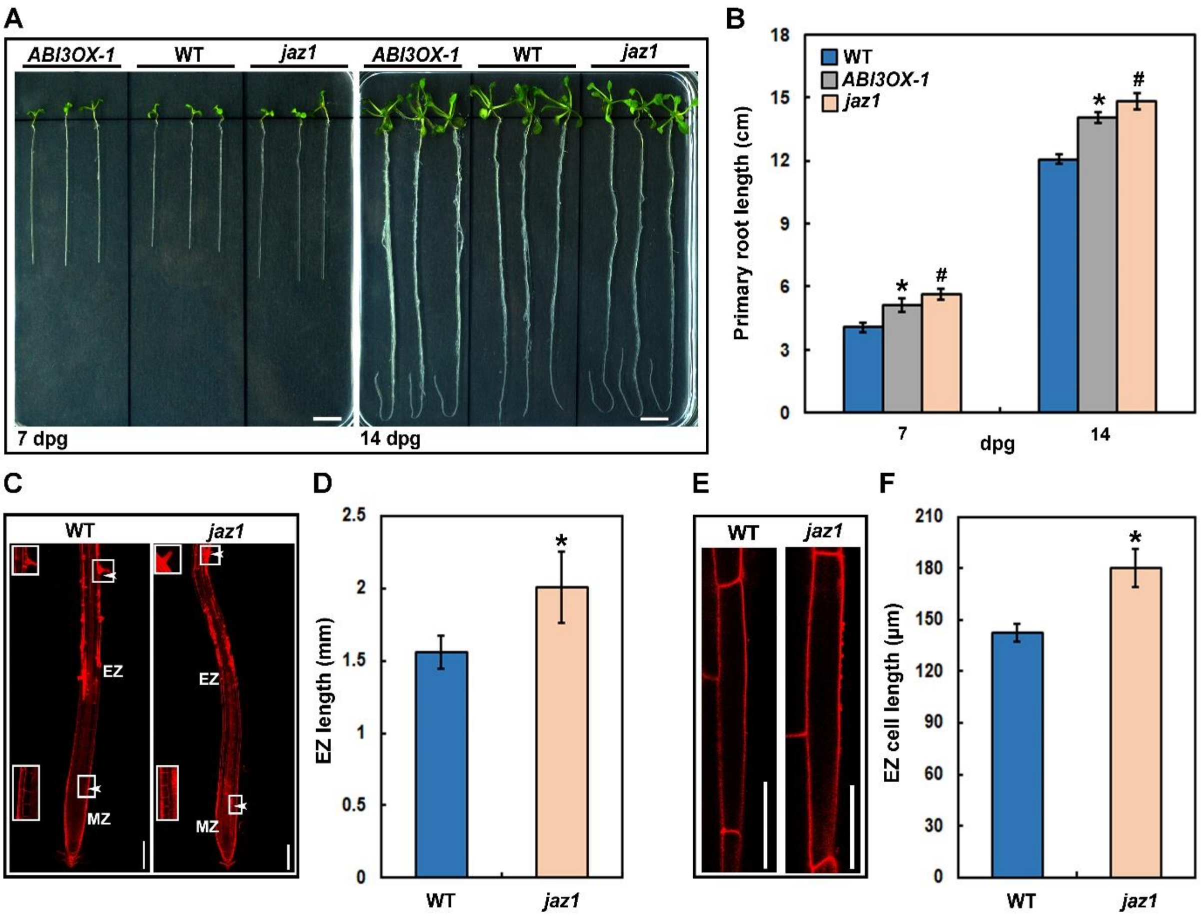


Figure S3

**Figure S3: *jaz1* mutant seedlings exhibit longer primary root growth phenotype**. **A.** Primary root growth of WT, *jaz1* and *ABI3OX-1* grown vertically on Murashige and Skoog (MS) medium for 7 and 14 days post germination (dpg). Scale bar: 1 cm. **B.** Primary root length measurement of WT, *jaz1* and *ABI3OX-1* over time. Data represent mean of three replicate experiments with n ≥ 20 seedlings for each replicate of each genotype with standard error of mean (SE). Anova two-factor with replication method was used to measure the variance, and the results with $P \leq 0.05$ considered to be significant. Variance between WT and *ABI3OX-1* is denoted by *, while variance between WT and *jaz1* is denoted by #. **C.** Longitudinal view of propidium iodide stained 7-day-old WT and *jaz1* primary root, where the elongation zone (EZ) is represented in between the two arrowheads. The initiation of first elongated root hair representing the differentiation zone, and the junction of the EZ and transition zone is shown in the inset of each primary root. MZ represents meristematic zone. Scale bar: 200 µm. **D.** Measurement of elongation zone length represented in (C) from 7-day-old WT and *jaz1* seedlings. Data represent means from three independent replicates (n ≥ 15) and error bars represent SE. Student's t-test with paired two-tailed distribution was used for statistical analysis, and $P \leq 0.05$ is denoted by *. **E.** Confocal images of epidermal root cells of the elongation zone (EZ) stained with propidium iodide from 7-day-old WT and *jaz1* seedlings. Scale bar: 50 µm. **F.** Measurement of epidermal root cells of elongation zone represented in (E) from 7-day-old WT and *jaz1* seedlings. Data represent means from three independent replicates (n ≥ 15) and error bars represent SE. Student's t-test with paired two-tailed distribution was used for statistical analysis, and $P \leq 0.05$ is denoted by *.

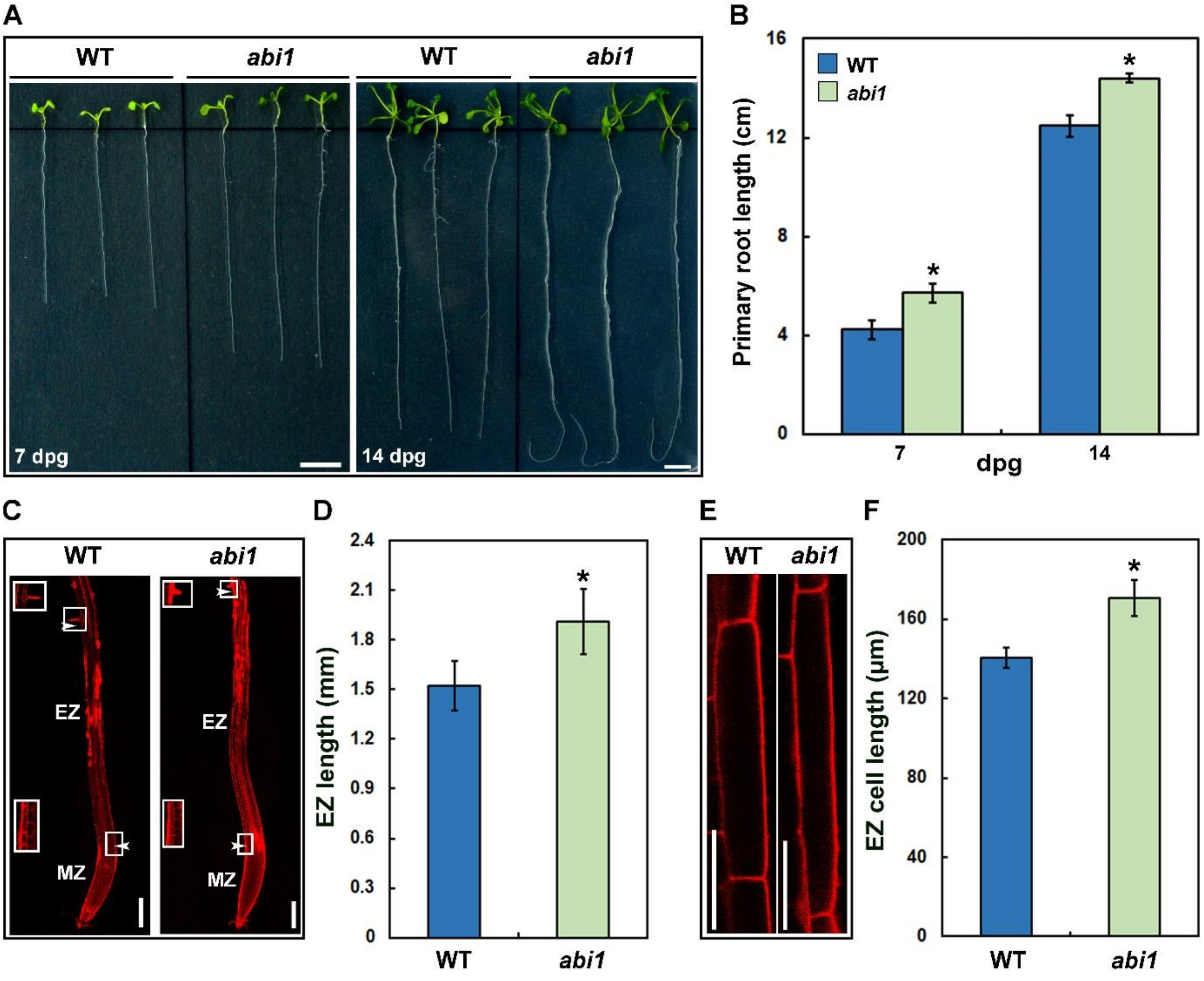


Figure S4

**Figure S4: *abi1* mutant seedlings exhibit longer primary root growth phenotype. A.** Primary root growth of WT and *abi1* grown vertically on Murashige and Skoog (MS) medium for 7 and 14 days post germination (dpg). Scale bar: 1 cm. **B.** Primary root length measurement of WT and *abi1* over time. Data represent the mean of three replicate experiments with n ≥ 20 seedlings for each replicate of each genotype with standard error of mean (SE). Student's t-test with paired two-tailed distribution was used for statistical analysis, and $P \leq 0.05$ is denoted by *. **C.** Longitudinal view of propidium iodide stained 7-day-old WT and *abi1* primary root, where the elongation zone (EZ) is represented in between the two arrowheads. The initiation of first elongated root hair representing the differentiation zone, and the junction of the EZ and transition zone is shown in the inset of each primary root. MZ represents meristematic zone. Scale bar: 200 µm. **D.** Measurement of elongation zone length represented in (C) from 7-day-old WT and *abi1* seedlings. Data represent means from three independent replicates (n ≥ 15) and error bars represent SE. Student's t-test with paired two-tailed distribution was used for statistical analysis, and $P \leq 0.05$ is denoted by *. **E.** Confocal images of epidermal root cells of the elongation zone (EZ) stained with propidium iodide from 7-day-old WT and *abi1* seedlings. Scale bar: 50 µm. **F.** Measurement of epidermal root cells of elongation zone represented in (E) from 7-day-old WT and *abi1* seedlings. Data represent means from three independent replicates (n ≥ 15) and error bars represent SE. Student's t-test with paired two-tailed distribution was used for statistical analysis, and $P \leq 0.05$ is denoted by *.

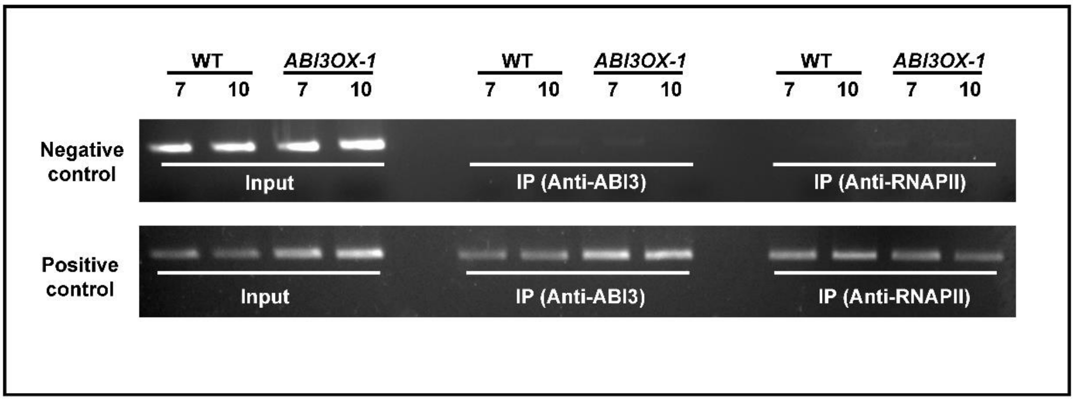


**Figure S5**

**Figure S5: ChIP-PCR for negative controls of ABI3 and RNAPII occupancy.** ChIP assay done using anti-ABI3 and anti-RNAPII antibody of 7- and 10-day-old root tissues of WT and *ABI3OX-1.* Agarose gel image showing PCR results using primers designed for regions that do not contain potential ABI3 and TATA box (top panel) and contain ABI3 binding sites and TATA boxes in the *JAZ1* locus (bottom panel). This data supports the specificity of results shown in Fig 5.

**Methods S1.** Detail description of the methods used in this study

**Floral dip method for transgenic generation**

Floral dip was done as described previously (Clough and Bent, 1998, Zhang et al., 2006). Overnight grown culture of *Agrobacterium tumefaciens* strain LBA4404 was resuspended in inoculation medium (Clough and Bent, 1998) (5% sucrose, 0.05% Silwet L77). The floral buds of 4-week-old WT (Col-0) and *abi3* were dipped in the inoculation medium and wrapped with plastic films to maintain high humidity for 16-24 hr (Zhang et al., 2006). The dipped plants were grown for 1 month at 22°C, 16 hr light/8 hr dark. The primary transformants were selected through appropriate antibiotic screening.

**RNA isolation for gene expression analyses**

Total RNA was isolated using TRIzol reagent (Thermo Fisher Scientific, Waltham, MA, USA) strictly as per manufacturers' protocol and cDNA was prepared from 5 µg of total RNA with RevertAid Reverse Transcriptase (Thermo Fisher Scientific, EP0441). The synthesized cDNA was then subjected for gene expression analyses by qPCR using gene specific primers (Table S1).

**Chromatin immunoprecipitation assay**

The isolated chromatin was incubated with anti-ABI3 (BioBharati Life Sciences, Kolkata West Bengal, India) (Bedi et al., 2016; Sengupta et al., 2020), anti-RNAPII (Abcam ab817), anti-H3K4me3 (Abcam ab8580), anti-H3K27me3 (Abcam ab195477) and anti-H3 (BioBharati Life Sciences, India) antibody overnight at 4°C. Protein-A/G agarose beads (BioBharati Life Sciences, India) were used to pull down the immune-complexes. qPCR with specific primer set (Table S1) was done to check the occupancy of ABI3 and RNAPII and histone modification levels. The data represent the mean of three independent biological replicates. For negative control, semi-quantitative ChIP-PCR was done with primers specific to *JAZ1* locus that contain no potential ABI3 or RNAPII binding sites.

**Real-time PCR and data analyses**

For quantitative real-time PCR analysis (qPCR), Dynamo Colour Flash SYBR Green qPCR Kit (Thermo Fisher Scientific, F416L) was used as per the manufacturer's instructions. For gene expression analyses diluted cDNA was used and *ELF1* was used as internal control. The expression level of each gene was determined using the formula $2^{-(Ct\ [sample]\ -\ Ct\ [internal\ control])}$

(Sengupta et al 2020, Bedi et al 2016, 2018, Mandal et al 2023). In the ChIP experiment, the $C_t$ values of the input samples were extrapolated to represent 100% since the input samples constituted 10% of the cell extract and the $C_t$ values of the IP (immune-precipitated DNA) samples were then plotted as a percentage of the input using the formula $100 * 2^{(Ct\,[Input] - Ct\,[IP])}$ (Sengupta et al 2020, Bedi et al 2016, 2018, Mandal et al 2023). For ChIP experiments with antibodies specific to histone modifications, all data was normalised to H3 levels. All PCR reactions were performed in triplicate, and the error bars indicate the standard error of mean (SEM). Statistical analysis was conducted using paired Student's t-test with a two-tailed distribution (for ChIP assay) and Anova two-factor with replication method (for gene expression analyses).

**Fluorometric *GUS* assay**

Leaves of 6-week old *Nicotiana benthamiana* or *Arabidopsis thaliana* (WT, *abi3* and *ABI3OX-1*) seedlings were infiltrated with *Agrobacterium tumefaciens* strain LBA4404 harbouring recombinant plasmid. For co-infiltration, *A. tumefaciens* containing different recombinant plasmids were mixed at a 1:1 ratio prior to infiltration. GUS measurements were obtained from at least 20 leaves ($n \geq 20$) on three biological replicates over three separate occasions. Error bar represents SE, and significant differences were calculated using a paired Student's t-test with two-tailed distribution. $P \leq 0.001$ was denoted by * on corresponding graphs.

**Histochemical *GUS* staining**

*Agrobacterium tumefaciens* strain GV3101 harbouring recombinant plasmid was infiltrated into leaves of 6-week-old *Nicotiana benthamiana* seedlings along with *P19* containing plasmid (Angel et al., 2011) to inhibit gene silencing. pCAMBIA 1304 and pKGWFS7 were used as positive and negative controls respectively. For co-infiltration, *A. tumefaciens* containing proJAZ1::GUS and 35S::ABI3 were mixed at a 1:1 ratio prior to infiltration. Infiltration was done in the abaxial side of leaves with 1 ml syringe (without needle). Plants were kept in dark for 12 hours and transferred in light for 24-36 hours. For histochemical GUS visualization, infiltrated leaves were incubated for 12 h at 37°C in the GUS-staining solution containing 0.1 M sodium-phosphate buffer (pH 7), 10 mM EDTA, 0.1% [v/v] triton X-100 and 1 mg/ml 5-bromo-4-chloro-3-indoxyl-β-D-glucuronide cyclohexyl ammonium salt (X-Gluc) (HiMedia Laboratories Pvt, India). Leaves were photographed after chlorophyll removal using 70% ethanol.

**Immunoblot and quantitative analyses**

Following separation of proteins (150 µg for ABI1 protein, 50 µg for PM $H^+$-ATPase, 100 µg for $pT^{947}$) in 8% SDS-PAGE, individual proteins were transferred to PVDF membrane in presence of transfer buffer. The membrane was blocked using 5% non-fat dried milk for 30 min and then incubated with specific antibodies overnight at 4°C. Polyclonal antibody against the phosphorylated Thr947 of AHA2 of was raised using the phosphorylated synthetic peptide CIETPSHYpTV, where pT represents phosphorylated threonine (Gene to Protein, India), as an antigen (BioBharati Life Sciences, India). Anti-ABI1 (Agrisera AS12 1861), anti-$H^+$-ATPase (Agrisera AS07 260) and anti-$pThr^{947}$ was used at a dilution of 1:3000 (TBS), 1:2500 (TBS) and 1:7500 (TBST) respectively. Thereafter membranes were incubated with secondary antibody, goat anti-Rabbit IgG, (BioBharati, India) for 1 hour. The membrane was developed using chemiluminescent substrate (SuperSignal® West Pico, Thermo Scientific) after adequate washes with wash buffer (50 mM Tris-Cl pH 7.4, 150 mM NaCl, 0.1% Tween 20). Western blot with anti-H3 antibody (BioBharati, Kolkata, India), was used as in internal control. Band intensities were quantified using ImageJ software. Each experiment was repeated for at least three independent biological replicates.

**Total protein isolation**

To detect ABI1 protein levels, 7-day old WT and *abi3* seedlings were treated with or without 50 µM ABA (HiMedia Laboratories Pvt, India) for 6 hours. The root samples were flash frozen in liquid $N_2$. The frozen samples were powdered using mortar-pestle followed by solubilization in extraction buffer (10 mM HEPEs, (pH 7.5), 100 mM NaCl, 1 mM EDTA, 10% glycerol, 0.5% Triton X-100, 1x Roche complete™ Protease Inhibitor Cocktail and 1 mM PMSF) (Kong et al., 2015). The samples were centrifuged at 10,000g and supernatant was collected. Protein concentration was estimated using Bradford reagent (Thermo Fisher Scientific).

To determine plasma membrane $H^+$-ATPase levels, 7 and 10-day old WT and *abi3* plant roots were flash frozen in liquid $N_2$. The frozen samples were powdered using mortar-pestle followed by solubilization in extraction buffer (25 mM Tris-HCl, pH 7.5, 150 mM NaCl, 1% Triton X-100, 1x Roche complete™ Protease Inhibitor Cocktail, 1 mM EDTA, 1 mM DTT and 0.5 mM PMSF) (Li et al., 2021). The samples were incubated on ice for 30 minutes, followed by centrifugation at 10,000g to discard the plant debris. The supernatant was recovered, and total protein was estimated using Bradford reagent (Thermo Fisher Scientific).

**Table S1: List of primers**

**A. For gene expression analysis**

| Accession number | Primer name | Sequence |
|---|---|---|
| AT1G19180 | JAZ1 FP | CTGATGTCAATGGAACTTTAGG |
| | JAZ1 RP | GTCATTGAATACAATCACTTGC |
| AT1G72450 | JAZ6 FP | GCTAAAGAAGCGAATCATGTTGC |
| | JAZ6 RP | CTAGCCACAGCCCTGTCTTTTC |
| AT2G34600 | JAZ7 FP | CATAGCTCGTTGGACGAATCAAG |
| | JAZ7 RP | CTTCATAGAGGCCTTTTGATAATG |
| AT1G30135 | JAZ8 FP | GATACAACCTCTGTTGTAGAATC |
| | JAZ8 RP | GGATTTGGAAGCTGATTATGATG |
| AT1G70700 | JAZ9 FP | CATATCTCCCGATAAGGCTCAAGC |
| | JAZ9 RP | CATAAGCCTCTCTTTGCGCTTC |
| AT3G24650 | ABI3 FP | CTTGAATGGGTCCAAACTA |
| | ABI3 RP | GGTCCGAGACAAAAGCTT |
| AT3G45140 | LOX2 FP | GATCTCATCAAAAGGGGGTTGGC |
| | LOX2 RP | AGTCATCTTGTGTTTTGAGGACAG |
| AT1G32640 | MYC2 FP | AGCAAACGGTAGAGAAGAGCCAC |
| | MYC2 RP | GACATATCTCCTCCACTAGCACTC |
| AT1G69530 | EXPA1 FP | ACGGACACTCTTACTTCAACCTAG |
| | EXPA1 RP | TAACTGCTTCTACTGTGAAGGTCTG |
| AT4G30190 | AHA2 FP | GCTGGTGTGATCTGGCTATACAG |
| | AHA2 RP | TCCCTAAGCCTAGCGATCTCAG |

**B. For cloning**

| Primer name | Sequence |
|---|---|
| ABI3 CDS FP | CACCATGAAAAGCTTGCATGTGGCGGC |
| ABI3 CDS RP | GCTCGGTTGTCTTACTTTAACCCCTCGTACT |
| proJAZ1 GW FP | CACCCACTCACTAACGCCGTTTAC |
| proJAZ1 GW RP | TTTGGTGCTGTTTTTTTTATCGGAT |

| | |
|---|---|
| ABI1 BamHI CDS FP | GGATCCATGGAGGAAGTATCTCCGGCGATC |
| ABI1 SalI CDS RP | GTCGACCAGATCCTCTTCTGAGATGAGTTTTT GTTCGTTCAAGGGTTTGCTCTTG |
| ARR1 BamHI CDS FP | GGGATCCATGAATTTGAAGAAACCGCGTG |
| ARR1 SalI CDS RP | GTCGACAACCGGAATGTTATCGATGGAG |

**C. For ChIP assay**

| Primer name | Sequence |
|---|---|
| JAZ1 ChIP FP | TTGACTTTGATGTATGACTTTTAGC |
| proJAZ1 GW RP | TTTGGTGCTGTTTTTTTTATCGGAT |
| JAZ1 CNC FP | CGGAGGATTATCAGGGAAATTGAAC |
| JAZ1 CNC RP | TCTTCTTCGATGCAGTCAAAGTAC |

**D. For mutant line screening**

| Primer name | Sequence |
|---|---|
| eGFP FP | TGGTGCCCATCCTGGTCGAGC |
| eGFP RP | GCTCGATGCGGTTCACCAGGG |
| Kanamycin FP | TCAGAAGAACTCGTCAAGAAGGCGA |
| Kanamycin RP | ATGATTGAACAAGATGGATTGCACGC |
| ABI1-038866 FP | CATTTAATGAAAGTCATCTTTATG |
| ABI1-038866 RP | CTCTGGTTGTGATCTATAAGATAG |
| JAZ1 gene specific FP | ATGTCGAGTTCTATGGAATGTTC |
| SALK LB-1.3 RP | ATTTTGCCGATTTCGGAAC |
